# Evolutionary conservation and innovations of RNA polymerase II transcription elongation factors

**DOI:** 10.64898/2026.01.11.698887

**Authors:** Alex M. Francette, Aakash Grover, Nathan Clark, Karen M. Arndt

**Author notes:** Authors who contributed equally to the work (Alex M. Francette, Aakash Grover).

## Abstract

In eukaryotes, transcription elongation factors (TEFs) associate with RNA Polymerase II (RNAPII) to facilitate gene expression and couple transcription to co-transcriptional processes, including chromatin regulation and RNA processing. To further our understanding of TEF biology, we developed a domain-centric analysis pipeline to perform a broad survey of ten TEF orthologs -- Paf1, Ctr9, Cdc73, Rtf1, Leo1, Spt4, Spt5, Spt6, Spn1, and Elf1 -- across the Tree of Life and analyze their evolutionary patterns in a structural context. We report evidence for all ten TEFs being present in the last eukaryotic common ancestor, indicating that mechanisms of TEF-mediated transcription regulation are both ancient and conserved. However, some early-diverging eukaryotic clades exhibit signs of altered TEF domain composition. A comparative phylogenetic analysis highlighted conserved regions of TEFs that are detected in both metazoans and fungi and other regions that appear clade-specific, detected only in metazoans. These observations, together with additional insights generated from evolutionary rate covariation analysis, shed light on under-characterized aspects of TEFs, including domains for which functions have yet to be dissected.

## Introduction

In all three domains of life – Bacteria, Archaea, and Eukaryota – genomic information is expressed in the form of RNA through the process of transcription. In eukaryotes, protein-coding genes are transcribed by the multi-subunit RNAPII and its associated factors. TEFs are one class of accessory factors that engage with active RNAPII to coordinate co-transcriptional events and promote RNAPII elongation through chromatin. Insights into TEF-mediated activity have been primarily derived from studies performed in eukaryotic model systems, such as fungi (*e.g. Saccharomyces cerevisiae*, *Schizosaccharomyces pombe*), invertebrates (*e.g., Drosophila melanogaster*, *Caenorhabditis elegans*), vertebrates (*e.g., Mus musculus*), and angiosperms (*e.g. Arabidopsis thaliana)* (1–4). From decades of study, several critical TEFs have been identified. Core TEFs, defined as those shared amongst major model organisms and visualized by cryo-EM of the activated transcription elongation complex called EC* include: Spt4(hSUPT4H) and Spt5(hSUPT5H) comprising DSIF, Spt6(hSUPT6H), Spn1(hIWS1), Elf1(hELOF1), and the five-subunit Polymerase Associated Factor 1 Complex or Paf1C – Paf1(hPAF1/PD2), Ctr9(hCTR9), Cdc73(hCDC73/Parafibromin), Rtf1(hRTF1), and Leo1(hLEO1) (4). For simplicity, these orthologs will be referred to by their *S. cerevisiae* protein designations.

The discovery and characterization of functionally conserved, often modular TEF domains have uncovered roles in promoting appropriate transcription. TEFs impact RNAPII elongation rate, are required for processive transcription *in vivo*, facilitate the transient removal and re-deposition of histones after RNAPII passage, and direct post-translational modifications to RNAPII and histones (1–6). However, the extent to which TEF sequences are conserved across diverse species remains unclear. In this study, we aimed to build an understanding of how the presence, organization, and conservation of these factors are distributed amongst diverse organisms to gain insights into the mechanisms of transcription regulation employed across the Tree of Life.

Modern analyses classify eukaryotes into several major lineages, many of which harbor divergent or poorly characterized transcriptional machinery. These lineages include the supergroups of Amorphea (*e.g*. Amoebozoa, Animalia, Fungi), Archaeplastida (Viridiplantae, Red Algae, Glaucophytes), Cryptista (*e.g*. *Cryptomonas*), CRuMs (Collodictyonid, Rigifilida, and *Mantamonas*), Discoba (*e.g*. *Trypanosoma, Euglena*), Haptista (*e.g*. *Choanocystis*), Metamonada (*e.g*. *Giardia, Trichomonas*), and TSAR (Telonemids, Stramenopiles, Alveolates, and Rhizaria; *e.g*. brown algae, *Plasmodium*, *Toxoplasma*) (7). The loosely defined “Excavate” groups of Discoba, Metamonada, and Malawimonadae and the orphan group of Ancyromonada are considered some of the more basal lineages of eukaryotes, placing them in an important position to understand the acquisition or loss of TEFs throughout evolution (7–9).

A limited number of TEFs are shared between prokaryotes and eukaryotes, and several studies have characterized TEF homologs in single-celled eukaryotes (10). Spt5 (NusG in bacteria) is the only TEF previously identified in all domains of life (11, 12). In Archaea and Eukaryota, Spt5 dimerizes with a small, globular protein, Spt4 (13), and Elf1 homologs also have been detected within both clades (10, 14). Spt6 shares structural homology with the bacterial transcription accessory factor Tex (11, 15–17). Recently, Paf1 and Spt6 have been characterized as deeply conserved in eukaryotes (18). However, the conservation of Spn1 and the remaining Paf1C subunits is less understood. Several investigations into eukaryotic TEFs outside of animals, fungi, and plants have been performed. Studies in *Trypanosoma brucei* found evidence for orthologs of Spt4, Spt5, Spt6, Ctr9, Cdc73, and Leo1 (19). Another study reported Elf1 orthologs in Apicomplexa species including the obligate intracellular parasite, *Toxoplasma gondii*, which are proposed to be functionally divergent (20). There are some indications that specific TEFs are not ubiquitous throughout Eukaryota, particularly in some parasites. A BLASTP search across 11 distantly related eukaryotes, including members of TSAR, Discoba, Metamonada, and Amoebozoa, failed to identify Paf1C subunits in Metamonads (*Trichomonas* and *Giardia*) (19). However, from the remaining species, Cdc73 was detected in all, and Ctr9 was found in all but *Plasmodium falciparum*. Paf1, Leo1, and Rtf1 were variably identified in this search. In another study, precipitation of RNAPII-associated factors in *Trypanosoma brucei* failed to identify Paf1, Rtf1, Spn1, or Elf1 (21). More recently, BLASTP and TBLASTN searches of Microsporidia species (an intracellular parasitic sister clade to fungi) variably identified homologs of Spt5, Spt6, and Paf1C (22). In essence, the landscape of elongation factor conservation is unclear, and it remains unknown whether the Last Eukaryotic Common Ancestor (LECA) had some or all these factors.

Seeking a deeper, unified analysis of their conservation and structural heterogeneity, we investigated the presence and co-occurrence of ten core TEFs in the proteomes of 304 broadly diverged species. Our analysis of TEF domain architecture suggests that the composition and general form of these TEFs were mostly defined in the LECA. However, we also identify multiple instances of apparent divergence in the composition of transcription elongation machinery. Furthermore, we compared the conservation of residues between orthologous proteins in fungal and metazoan proteomes to identify specific residues and regions of common and clade-specific importance. These findings home in on recently identified protein-protein interaction interfaces and identify additional regions of potential functional importance. Together, our analysis provides both broad and in-depth insights into the conservation and function of factors fundamental to eukaryotic transcription.

## Results

### Paf1C and Spn1 are only detected in eukaryotes

Since several TEFs are structurally modular and contain more than one identifiable domain (**Fig. 1A**), some orthologous proteins may not share the exact structural organization of known TEFs. Previous investigations searching for TEF orthologs along the entire protein sequence may have been biased against those with divergent architectures. Therefore, to minimize this bias, we took a domain-centered approach to identify orthologs of Paf1, Ctr9, Cdc73, Rtf1, Leo1, Spt4, Spt5, Spt6, Spn1, and Elf1 across a diversity of bacterial (n = 37), archaeal (n = 40), and eukaryotic (n = 227) proteomes (**Fig. 1A** and **Fig. S1**) (**Methods**). For this work, a “domain” was operationally defined as a functional unit of a protein supported by structural, genetic, and/or biochemical evidence, irrespective of its ability to form a stable fold. Rather than relying solely on established domain families such as those in the PFAM database (23), we adopted a customized strategy to capture a wider range of putative orthologs. We searched proteomes from the EukProt and GTDB databases for each protein of interest using both BLAST, with whole protein sequences as templates, and custom hidden Markov models (HMMs) built with each domain (see **Methods**). The resulting hits were then individually inspected and filtered to remove spurious identifications, a process guided by a combined analysis of multiple sequence alignments (MSAs), gene tree reconstructions, and further searches using both PFAM-provided and custom HMMs. This methodology, informed by extensive structural and functional studies of the target proteins, was designed to be a sensitive survey that captures orthologs that may have diverged significantly over time. All domain-specific searches outperformed BLAST, allowing expansion of the range of identified orthologs (**Fig. S2-S3**). We note that our ability to detect a domain using this pipeline in any given proteome is limited by several factors including: 1) the source of the predicted proteome either from transcriptomic profiling or genomic sequence (**Fig. S4A**); 2) the completeness of the predicted proteome (**Fig. S4B, C)**; 3) the amount of information in the HMM for the domain (**Fig. S4D)**; and 4) the sequence divergence of the target domain in a given proteome.

**Figure 1.**
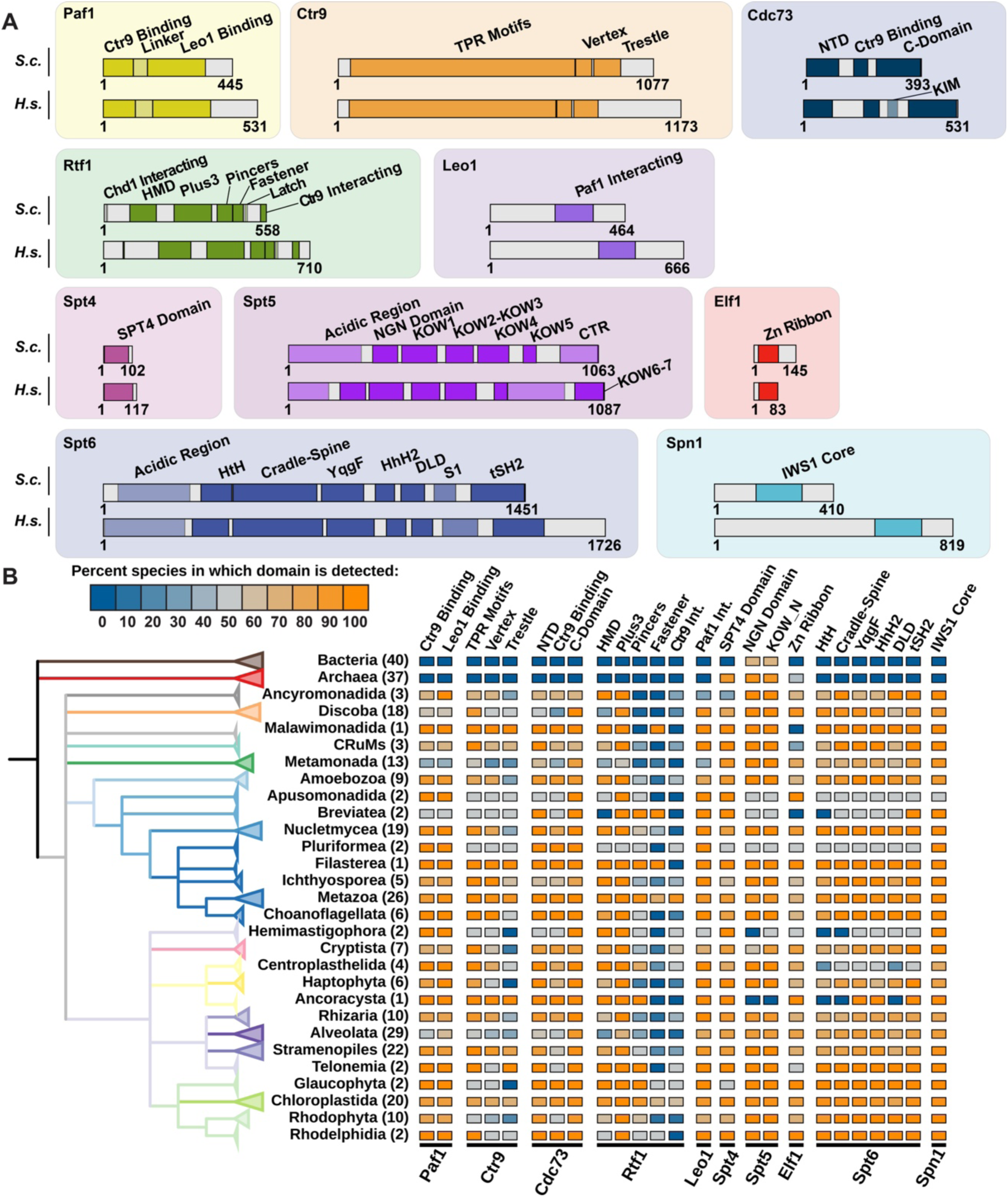
TEF domains are broadly detected across Eukaryotes. (A) Architecture of core TEFs, highlighting regions with verified functions. Domains utilized for homolog searches are in dark shading. Additional domains too small or unstructured to be confidently used in homolog searches are highlighted with light shading. (B) Distribution of TEF domain detection across the Tree of Life. The number of species analyzed in each clade is shown in parentheses. Each column represents the percentage of detection across clades for the indicated domain. Unless otherwise mentioned, the Fastener indicated in Rtf1 corresponds to the RNAPII-fastener as defined in (28). TPR - Tetratricopeptide Repeat; NTD – N-Terminal Domain; HMD – Histone Modification Domain; NGN – NusG N-terminal; KOW - Kyrpides-Ouzounis-Woese; CTR – C-Terminal Repeat; HtH – Helix-Turn-Helix; HhH2 – double-helix-hairpin-helix; DLD – Death-Like Domain; tSH2 – Tandem Src Homology 2.

Using this domain-centric pipeline, we recapitulate observations made by previous studies focused on the evolution of transcription factors (11, 24). Spt5, identified by the presence of the NGN domain and at least one KOW domain, is the only universally conserved TEF across kingdoms (**Fig. 1B** and **Fig. S5**). Spt4 orthologs, identified by their zinc-finger (hereafter, the SPT4 Domain; **Fig. 1A**), are detected in most archaea and eukaryotes. Elf1 orthologs, characterized by the presence of an Elf1 zinc-finger (**Fig. 1A**), are detected in most of the Asgardarchaeota, some non-Asgard archaeal species, and most eukaryotes.

Spt6 domains, Paf1C domains, and the Spn1 IWS1 domain were only detected in eukaryotes. Given the structural similarity between the Spt6 and Tex core domains, we expected that our search would identify Tex homologs from prokaryotic proteomes in addition to Spt6 homologs from eukaryotes. However, using our HMMs, we were only able to detect Spt6 homologs in eukaryotes (**Fig. 1B** and **Fig. S5**). These observations indicate that while extant Tex and the core of Spt6 derived from a common ancestor, over the course of evolution, the core domains in Spt6 have diverged from Tex at the sequence level. Our inability to detect Paf1C domains or the Spn1 IWS1 domain in any prokaryotic lineage prompted us to ask if we could detect these domains in a larger set of 218 Asgardarchaeota proteomes (**Fig. S4C**), the closest known archaeal clade to eukaryotic species (25). While we were able to detect Spt5, Spt4, and Elf1 orthologs (**Fig. S4E**), we were unable to detect any proteins that contained the IWS1 domain or Paf1C domains, even in this targeted search. Thus, we propose that Paf1C and Spn1 are eukaryotic innovations.

### Some Paf1C domains are not detected in sub-clades of Discoba, Metamonada, and Alveolata

Most TEF domains were detected in basal eukaryotes, suggesting that these TEFs were present in the LECA (**Fig. S5**). However, while some TEFs, like Spt4, Spt5, Spt6, and Spn1, were almost universally detected across eukaryotic clades, the detection of some domains in Paf1C and the Elf1 domain was less consistent (**Fig. S5**). The under-detection of the Elf1 domain did not display any clade-specific pattern (**Fig. S5** and **Fig. S6A**). In contrast, several Paf1C domains were not detected in specific discobid, metamonad, and alveolate sub-clades (**Fig. S6A** and **Fig. 2A**). We observed two patterns of domain under-detection. First, in some sub-clades, we were unable to detect any domain corresponding to the proteins of interest (**Fig. 2A**). For example, we were unable to detect the Ctr9-binding and the Leo1-binding domains of Paf1 in Kinetoplastea and Parabasalia, sub-clades of Discoba (gray box 1) and Metamonada (gray box 2), respectively. Second, in some sub-clades, we detected at least one but not all domains corresponding to the protein (**Fig. 2A**). For example, in the Kinetoplastea and Diplonemea sub-clades of Discoba (gray boxes 1 and 2, respectively), we detected proteins containing the Rtf1 Plus3 domain but not the Rtf1 HMD.

**Figure 2.**
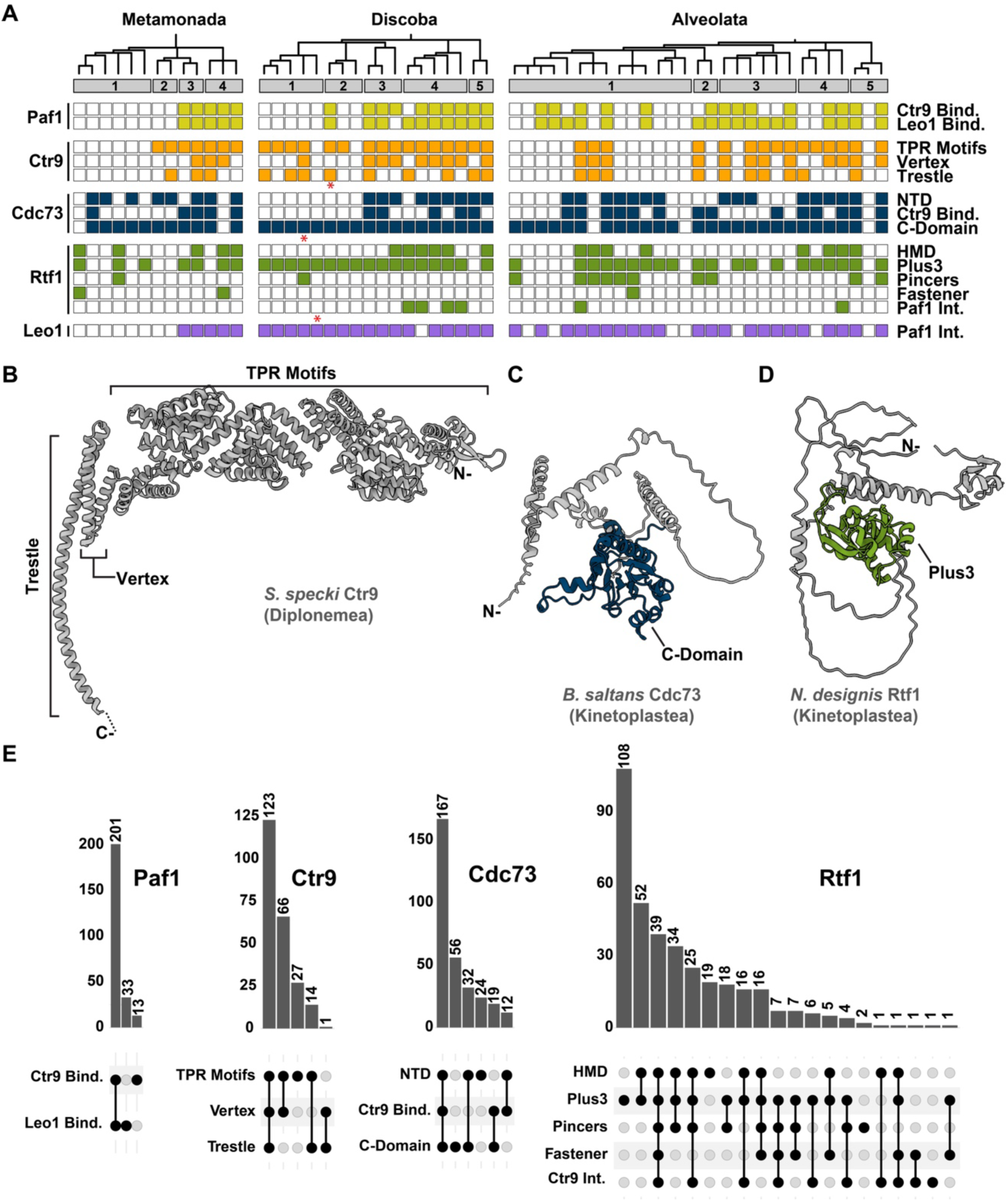
Paf1C domains are variably detected in discobids, metamonads, and alveolates. (A) Binary presence-absence heatmaps for Paf1C domain detection in Discoba, Metamonada, and Alveolata. Dendrogram on top represents phylogenetic relationship between species, as shown in Fig. S5. Gray bars represent sub-clades within the three clades. Metamonda: 1) BF clade (Fornicata and Barthelona), 2) Parabasalia, 3) Anaeramoebidae, and 4) Preaxostyla. Discoba: 1) Kinetoplastea, 2) Diplonemea, 3) Euglenida, 4) Heterolobosea, 5) Jakobida. Alveolata: 1) Apicomplexa, 2) Colpodellida, 3) DP clade (Dinoflagellata and Perkinsea), 4) Ciliophora, 5) Colponemidae. (B-D) AlphaFold3 predicted structures of Paf1C orthologs from indicated clades (marked by red asterisks in panel (A). (B) Model of *S. specki* (Diplonemea) Ctr9, indicating the presence of the Vertex in this clade. (C) Model of *B. saltans* (Kinetoplastea) Cdc73. The NTD is not detectable in proteomes from this clade. (D) Model of *N. designis* (Kinetoplastea) Rtf1. The HMD is not detected in proteomes from this clade. (E) UpSet plots depicting the coincidence of domain detection in homologs of indicated proteins as determined by HMMER hmmscan (97) using custom HMMs. Scan domain eValue (--domE) threshold set to 10^-3^.

Our inability to detect some domains in these clades can be attributed to the overall lower completeness of their predicted proteomes (**Fig. S6B**), a technical limitation of our approach, or a legitimate evolutionary loss event. We therefore used alternative approaches to search for Paf1C domains in clades in which our HMM search was unable to find a given domain (**Supplementary Table 1**). Alternative approaches included FoldSeek to search the AlphaFold predicted structure database of UniProt proteomes, manual inspection of AlphaFold3 predicted structures of hits obtained in our search, and PSI-BLAST searches of NCBI proteomes (see **Supplementary Methods**). In our HMM-based search, the Rtf1 HMD and Cdc73 NTD were undetectable in the Colpodellida sub-clade of alveolates (gray box 2). In contrast, FoldSeek was able to detect these domains in this sub-clade (**Supplementary Table 1**). Similarly, the Ctr9 Vertex was undetectable in the Parabasalia and Diplonemea sub-clades by our search. However, AlphaFold3 structure predictions of Ctr9 orthologs identified by our HMM and belonging to this sub-clade indicate that this domain is indeed present (**Fig. 2B, Fig. S7A, and Supplementary Table 1**).

Manual inspection of AlphaFold3 predicted structures of Cdc73 homologs in Kinetoplastea and Diplonemea indicate that while the Cdc73 C-Domain is detected in these sub-clades, these proteins lack a detectable NTD (example shown in **Fig. 2C**). Similarly, the Rtf1 HMD was not evident in predicted structures of Rtf1 homologs in these two clades, while the Plus3 domain was readily detectable (example shown in **Fig. 2D**). These observations are consistent with our HMM search which was unable to detect the Rtf1 HMD and Cdc73 NTD in Kinetoplastea and Diplonemea sub-clades (**Supplementary Table 1** and **Fig. 2A**). These approaches were also unable to detect the Paf1 Leo1-binding domain or Leo1 Paf1-binding domain in Parabasalia and Fornicata sub-clades of Metamonada (gray boxes 1 and 2) or the Rtf1 HMD in the Dinoflagellata sub-clade of alveolates (gray box 3). Altogether, these results indicate that select sub-clades of unicellular eukaryotes may have lost some Paf1C domains or the sequences of these domains are too diverged to be detected by the employed methods.

### Multi-domain architectures of TEFs are broadly conserved

A striking feature of TEFs like Rtf1, Spt5, Paf1, Cdc73, Ctr9, and Spt6 is their modular, multi-domain architecture. To determine if domain architecture is conserved for these six TEFs, we collected the identified homologs for each TEF and asked which of the domains are detected in each homolog by HMM scans. We then represented these data as UpSet plots (**Fig. 2E** and **Fig. S7B-C**). We find that, in general, for all multi-domain TEFs except Rtf1, the most commonly detected domain architecture is one in which all known domains are detected. In the case of Rtf1 (**Fig. 2E**), the most common domain architecture is the one in which only the Plus3 domain is detected. The next most common architecture for identified Rtf1 homologs is the co-occurrence of the HMD and Plus3 domain. The lack of detection of other Rtf1 domains (Pincers, Fastener, and the Ctr9-interacting region) in the identified homologs might relate to the fact that the HMMs of these domains have relatively lower information content than the two other Rtf1 domains (**Fig. S4D**).

### Conservation score analysis pinpoints functionally conserved residues in TEFs

A slow rate of amino acid substitution, *i.e.* a high degree of conservation, in orthologous proteins can suggest the functional importance of specific residues. Using our dataset of TEF orthologs, we calculated conservation scores for residues along human TEF sequences (see **Supplementary Methods**). As expected, previously characterized functional regions are generally more conserved than other regions, though not uniformly so (**Fig. 3A** and **Fig. S8A**).

**Figure 3.**
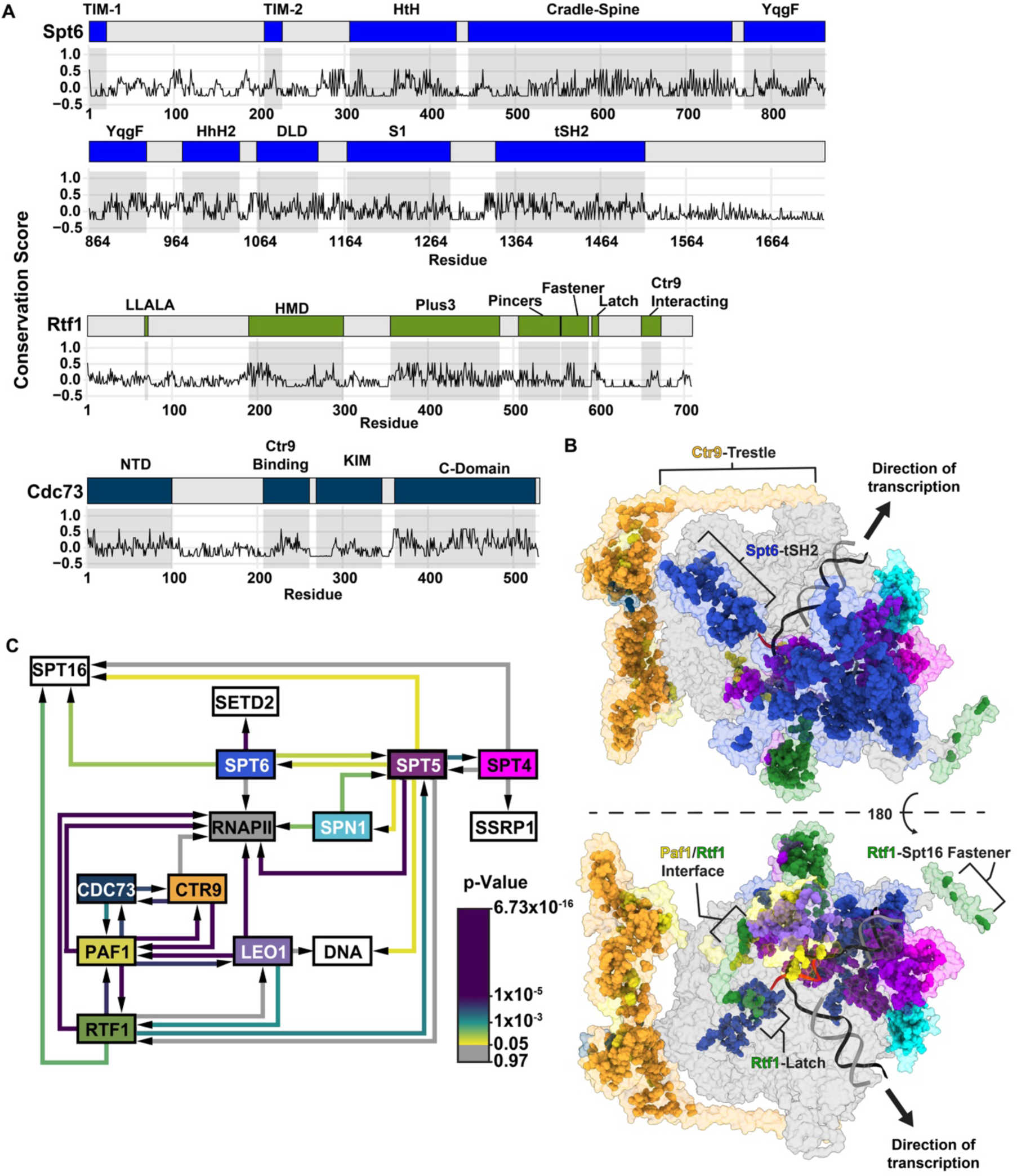
Conserved regions in TEFs. (A) Conservation scores (-log10(RER + 0.1)) of Spt6, Rtf1, and Cdc73 across each residue along the *H. sapiens* homolog. The domain map of each protein is stacked on top of a line plot showing the conservation score of each residue. Gray boxes in the line plot highlight domain boundaries. (B) Structure of transcription elongation complex (PDB: 9EH2) with atoms of conserved residues (conservation score > 0.28) shown as spheres. Nucleic acids are represented as ribbons and RNAPII surface is shown in gray. SPT16, SSRP1, and SETD2 are hidden for clarity. The structure was prepared in the absence of human Elf1 homolog, ELOF1. (C) Network of pairwise interactions between TEFs and other components of the transcription elongation complex. The average conservation scores of surface-exposed residues within the interfaces (TEF residues within 4 Å of the interacting molecule and solvent-accessible surface area > 50 Å^2^) were compared to all surface-exposed residues in the protein by Signed Rank Test to determine if residues at a given interface have a conservation score significantly higher than those of other non-buried residues. Arrows originate from a TEF towards its interactor. Edge colors indicate p-value. Since conservation scores were not calculated for any RNAPII subunit, SPT16, SSRP1 or SETD2, no edges emerge from these nodes.

Mapping conserved residues (conservation score > 0.28) onto a recent structure of the human transcription elongation complex (PDB: 9EH2, lacking Elf1) (26) identifies the locations of inter- and intra-protein contacts (**Fig. 3B** and **Fig. S8B**), selectively highlighting regions of known functional importance. For example, core residues stabilizing structured domains, such as the Spt6 tSH2 domain (**Fig. 3B**), are markedly more conserved (27). Regions with defined regulatory roles are also highly conserved, such as the Rtf1 Latch, which stimulates RNAPII elongation rate (**Figs. 3A-B** and **Fig. S8A**) (28). In contrast, structurally conserved regions that lack strongly conserved residues likely serve primarily architectural roles. For example, few residues in the Ctr9 trestle exhibit particularly high conservation (**Fig. 3B** and **Fig. S8A**). Despite this observation, the Ctr9 trestle is detected by the HMM scan in 60% (138/231) of CTR9 homologs spread broadly across the Tree of Life (**Fig. 2E and Fig. S5**).

To systematically describe how macromolecular interfaces dictate evolutionary patterns in TEFs, we examined every binary interaction resolved in the 9EH2 structure (**Fig. 3C** and **Fig. S8C**) (26). We assessed whether surface-exposed residues within 4 Å of other proteins or DNA in this static view exhibit a higher-than-average conservation score relative to other surface-exposed residues of each TEF. We find that Paf1 is not only a central hub for interfaces with other Paf1C subunits, but that these interfaces are all significantly enriched for conserved residues from both sides of the interaction (**Fig. 3C**). For example, our analysis indicates specific residues at the interface between the Rtf1 Fastener and Paf1 Linker are more conserved relative to other structurally resolved residues in these domains (**Fig. 3B** and **3C**). Likewise, the residues of Paf1, Rtf1, Leo1, and Spt5 proximal to RNAPII are highly conserved (**Fig. 3C**).

Importantly, less conserved residues do not imply a lack of functional importance when there is positive selection or relaxed selective constraints such as the case with intrinsically disordered charge blocks in the Ctr9 C-terminal tail (29). Spt4, a small, globular protein with a deeply conserved interaction with Spt5 (13), provides another example. Given the compact nature of Spt4, much of its sequence is comprised of nearly invariant residues like the cysteine residues coordinating Zn^2+^ in its Zn-finger motif (**Fig. S8A-B**) (30). Since conservation score is normalized over the length of each protein, the conservation of these residues may drive down the relative conservation of residues at the Spt4/Spt5 interface. In essence, when most residues are extremely conserved, few seem more conserved than average. Together, this analysis indicates that the interfaces between Paf1C subunits and several points of attachment to RNAPII are conserved across species.

### Conserved residues in fungi and metazoans suggest regions with preserved functions

While *S. cerevisiae* is a prominent model for the study of eukaryotic transcription, the common ancestors of fungi and metazoans diverged approximately one billion years ago (31). Therefore, an important objective is to understand which TEF features are shared between fungi and humans and which now differ. We reasoned that if we compared the independent patterns of evolution of TEF orthologs in metazoan- and fungal-specific proteome databases, we could identify residues and structural elements that are concordantly conserved across both clades. Alternatively, we expected to identify residues and structural features that are more conserved in either clade. These regions would likely represent functional elements unique to or selectively lost from one clade.

To this end, we examined RefSeq metazoan and fungal proteome databases (32) for TEF orthologs. While our detection of TEFs in basal eukaryotes is variable (**Fig. 1B)**, the divergence of fungi, such as *S. cerevisiae,* from metazoans, including *H. sapiens,* occurred comparatively recently leading to a high frequency of detection for all factors using a standard BLAST search (**Fig. S9A**). Next, we calculated the conservation scores for fungal and metazoan orthologs and mapped *S. cerevisiae* TEF residues to *H. sapiens* sequences (**Fig. 4A**). We then examined patterns of TEF evolution in the context of the *H. sapiens* AlphaFold structural predictions (**Fig. 4B-D**, **Fig. S9B-D**).

**Figure 4.**
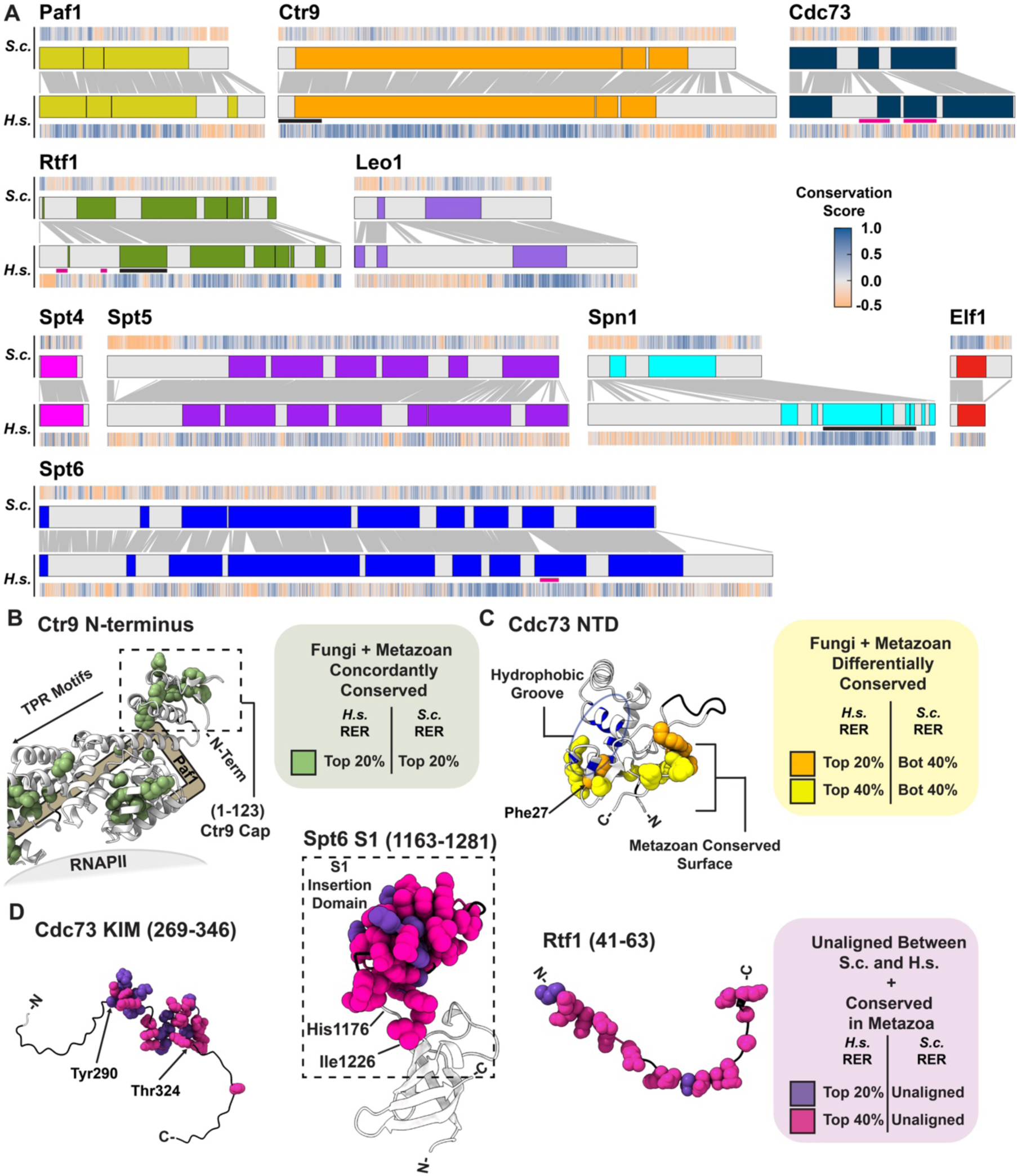
Similarities and divergences between metazoan and fungal TEF homologs highlight broad and clade-specific sites of potential functional importance. (A) Pairwise residue alignment (gray lines) of indicated proteins mapped on the linear domain maps of *S. cerevisiae* (top) and *H. sapiens* (bottom) homologs. Domains are as indicated in Fig. 3A and **Fig. S8A**. Heatmaps represent the conservation score of each residue for the indicated homolog. For *S. cerevisiae* and *H. sapiens*, conservation score was calculated using the alignment and gene trees built from fungal and metazoan homologs, respectively. Black and pink lines below the domain maps indicate regions that have been highlighted in panels B-D and **Fig. S9B-D**. Black lines indicate conserved regions in both fungi and metazoan homologs. Pink lines indicate conserved regions in metazoans that are not mappable in the fungal homologs. (B-D) AlphaFold2 predicted structures of human TEFs, highlighting different classes of conserved residues. Residues unaligned between the *S. cerevisiae* and *H. sapiens* orthologs and are not well conserved have been colored black. (B) Concordantly conserved residues (top 20% conserved residues in metazoan and fungal homologs) in the N-terminus of Ctr9. (C) Differentially conserved residues (top 20-40% conserved residues in metazoan homologs and bottom 40% conserved residues in fungal homologs) in Cdc73 NTD. Phe27 and residues labelled in blue form a hydrophobic surface groove that was previously suggested to be a potential binding surface (37). Residues in Cdc73 NTD selectively conserved in metazoans include Lys15, Lys16, Phe27, Asp109, Arg110, Ser111, Ala 112, Pro113, and Glu115. (D) Top 20-40% conserved residues in metazoan homologs that could not be aligned to the *S. cerevisiae* homolog. Conserved residues are highlighted in Cdc73 (left), an insertion within the S1 domain of Spt6 (middle), and near the Rtf1 N-terminus (right).

We first identified residues occupying homologous positions in the primary structures of TEFs that are amongst the most conserved in both fungi and metazoans (top 20%) (**Fig. 4B**, and **Fig. S9B**). These residues are expected to confer an evolutionary disadvantage when mutated in either *H. sapiens* or *S. cerevisiae* and thus likely mediate a shared role in Opisthokonts, which include both fungi and metazoans. As anticipated, we find core residues deep within protein folds, such as the IWS1 core domain in Spn1, in this class (**Fig. S9B, left**). However, this conservation in Spn1 extends C-terminally from the structured core into a space proximal to the sites where Elf1 and the Spt5 NGN bind to elongating RNAPII. This stretch was recently shown by cryo-EM to be directly interacting with human Elf1, Spt5 NGN, and the Rpb2 subunit of RNAPII (5, 6). We also identify residues extending from the Rtf1 HMD region that fasten to the histone chaperone Spt16 (26, 33) to be highly conserved in both clades (**Fig. S9B**).

The Ctr9 TPR motifs scaffold Paf1C assembly, mediating extensive interactions with Paf1 and Cdc73 (34–36). Interestingly, the N-terminal cap of Ctr9, consisting of two β-strands, a short α-helix, and the first TPR motif, is enriched with residues highly conserved in both fungi and metazoans (**Fig. 4B**). While yeast Ctr9 contains a 16 residue N-terminal tail that interacts with Paf1 and is important for Paf1-Ctr9 subcomplex assembly (34), there is no such extension in metazoan Ctr9. This cluster of conserved, solvent-facing residues in the Ctr9 cap is consistent with an additional role as a potential platform for interactors outside of Paf1C. Together, this analysis provides a global view of the conserved architectural cores and interaction surfaces maintained across evolutionary timescales.

### Differential conservation score analysis uncovers putative interfaces selectively conserved in Metazoa but not in Fungi

To explore how TEF biology may differ between fungi and metazoans, we investigated the possibility that structurally homologous regions in TEFs may be repurposed between clades. For example, if a purely architectural sequence acquires a new function as the site of an intermolecular interface, we would expect the identity of the residues involved in that specific interaction to change less over time. Therefore, we searched for residues that are slow to change in metazoans while rapidly changing in fungi. We categorized residues as differentially conserved if they are amongst the highest 20% or 40% conserved in *H. sapiens* and the lowest 40% conserved in *S. cerevisiae* (**Fig. 4C** and **Fig. S9C**).

This analysis identified a highly localized cluster of residues that are variable in fungi but remain strongly conserved in human Cdc73 (**Fig. 4C**). These residues are almost entirely distinct from a previously reported hydrophobic groove in the NTD speculated to be an interaction interface (37). Similarly, we note that a region of the Paf1 linker, between its known interacting sites with Rtf1 and Leo1, contains several differentially conserved residues, suggesting an uncharacterized role in metazoans (**Fig. S9C, left**). We additionally observe that the solvent-exposed surface of Ctr9 TPR motifs harbors two sites with differentially conserved residues, one of which lies in close proximity to the binding site of WDR61, a member of a surveillance complex that associates with mammalian but not yeast Paf1C (**Fig. S9C**, right) (38). The other site contains residues within TPR repeats 2-8 and lies just downstream of the N-terminal cap. These data highlight currently under-characterized regions in TEFs that might drive metazoan-specific functions or potentially mediate ancestral functions that have been lost in fungi.

### Sites of high conservation within regions unaligned between *H. sapiens* and *S. cerevisiae* TEFs

To survey core TEFs for potential functional features not observed in *S. cerevisiae*, we examined *H. sapiens* TEF sequences that failed to align to their *S. cerevisiae* counterparts for regions of high conservation (conservation score in highest 40%). With respect to Paf1C, we find that the Cyclin-K Interacting Motif (KIM) of Cdc73 (residues Tyr290-Thr324), recently shown to directly interact with CDK12/13 (39), is selectively identified out of the larger disordered Cdc73 linker to be highly conserved (**Fig. 4D**, left). As another example, the linker sequence between the NTD and Ctr9-binding domains in Cdc73 contains clusters of conserved residues in metazoans (**Fig. S9D**, left). Although some of these regions have defined functions as nuclear/nucleolar localization signals (residues 125-139 and 192-194 respectively) (40, 41), the region between Arg147 and Glu180 is additionally enriched for conserved residues (**Fig. S9D**, left). These residues overlap with Cdc73 residues 128-227, a site previously found to facilitate the association of Cdc73 with the H3 K9 methyltransferase, SUV39H1 (42). We also find two sites in human Rtf1 stand out as hotspots of conservation in areas that do not align to *S. cerevisiae* (**Fig. 4D**, right and **Fig. S9D**, right). One spans residues 41-63 immediately preceding the LLALA box (**Fig. 4D**), a site required for interaction between Rtf1 and homologs of the chromatin remodeler Chd1 (43). The other site (residues 149-162) lies between the Rtf1 LLALA box and the HMD (**Fig. S9D**, right). We predict these regions to contribute towards important functions of Paf1C in metazoans.

Intriguingly, whereas yeast and most other clades have a contiguous S1 domain fold in Spt6, the human protein exhibits a bipartite architecture bearing a 50-residue insertion, which we term the S1 Insertion Domain (SID) (**Fig. 4D**, middle). This region was recently resolved in a cryo-EM structure of a human transcription elongation complex (26) and had previously been noted in a comparison of S1-domain containing proteins as a peculiarity in the *Caenorhabditis elegans* Spt6 ortholog, EMB-5, which is absent in other S1 proteins (44). The SID is predicted to adopt a C2HC zinc-finger fold (**Fig. S10A**) and is juxtaposed to the RNA-exit channel in the RNAPII elongation complex (26, 28). However, the SID has an electronegative surface likely incompatible with RNA binding (**Fig. S10B**). Searches for similar inserts across eukaryotic S1 domains revealed the SID region to be ubiquitous in metazoans and variably detected in choanoflagellates (**Fig. S10C-D**), a sister group to Metazoa (45). Other eukaryotic species show no such insertion, excepting outliers amongst Alveolata and Metamonada (**Fig. S10C-D**). Thus, we conclude that the SID is not a universal feature of Spt6 but potentially originated prior to the divergence between choanoflagellates and metazoans.

### Support for a functional Ctr9-interacting Motif in human Rtf1

Unlike in available metazoan elongation complex structures, a C-terminal region of fungal Rtf1, which we term the Hook, can be resolved to associate with Ctr9 as a key attachment point to Paf1C (46–48). Consistent with these observations, metazoan Rtf1 is biochemically dissociable from Paf1C while fungal Rtf1 is a more stable member of Paf1C (49–52). Nonetheless, it remains unclear whether human Rtf1 directly engages with Ctr9. Near the C-terminus of human Rtf1, we find a stretch of conserved residues (655–674) that failed to align to the *S. cerevisiae* Rtf1 Hook region but aligned to the presumed Hook of other fungal Rtf1 orthologs (**Fig. 4A**, **Fig. S11A-B**). These residues in human Rtf1 have been previously reported as homologous to the fungal Ctr9-interacting region (47). Indeed, deleting the C-terminal residues from human Rtf1 containing this region (604–710) leads to reduced immunoprecipitation of other Paf1C subunits (53). These observations led us to examine the possibility that the human Rtf1 Hook interacts with Ctr9 using the same binding modality as yeast.

AlphaFold3 predictions suggest both human and yeast Rtf1 consistently fold onto the same groove of Ctr9 (**Figs. S11C-E**), albeit with the human pair having a slightly lower confidence. Furthermore, Ctr9 from both *S. cerevisiae* and *H.* sapiens bears a lipophilic pocket that appears biochemically compatible with the lipophilic face of the Rtf1 Hook (**Figs. S11F-G**). These data support a model where Rtf1 in metazoans, including humans, utilizes a linear motif to bind Ctr9, but this interaction may not be strong enough to prevent dissociation of Rtf1 from Paf1C.

### Evolutionary rate co-variation (ERC) landscape of TEFs

Our evolutionary analysis of TEFs has identified several domains and residues that are candidates for functional characterization in future studies. However, previous work indicates that co-variation of whole-protein evolutionary rates between genes is another powerful predictor of functional interaction amongst proteins (54, 55). Thus, we utilized a dataset generated from 343 yeast species to identify genes whose whole-protein evolutionary rates co-vary with TEFs. As done previously, we standardized the ERC values for each TEF by calculating Z-scores (56). For this analysis, we considered an arbitrary Z-score cut-off of 3.5 to identify genes that have a “high” ERC with TEFs. We then visualized genes that have a high ERC with the ten candidate TEFs as a network (**Fig. 5**). Consistent with their known physical and functional association, TEFs shared a high ERC with each other (black nodes connected with black edges). The interconnectedness of nodes in this network indicates that several genes share a high ERC with the same TEFs, resulting in a clustering coefficient much higher than expected from randomly generated networks of similar size (**Fig. S12A**). These genes include those that encode core RNAPII subunits and transcription initiation factors.

**Figure 5.**
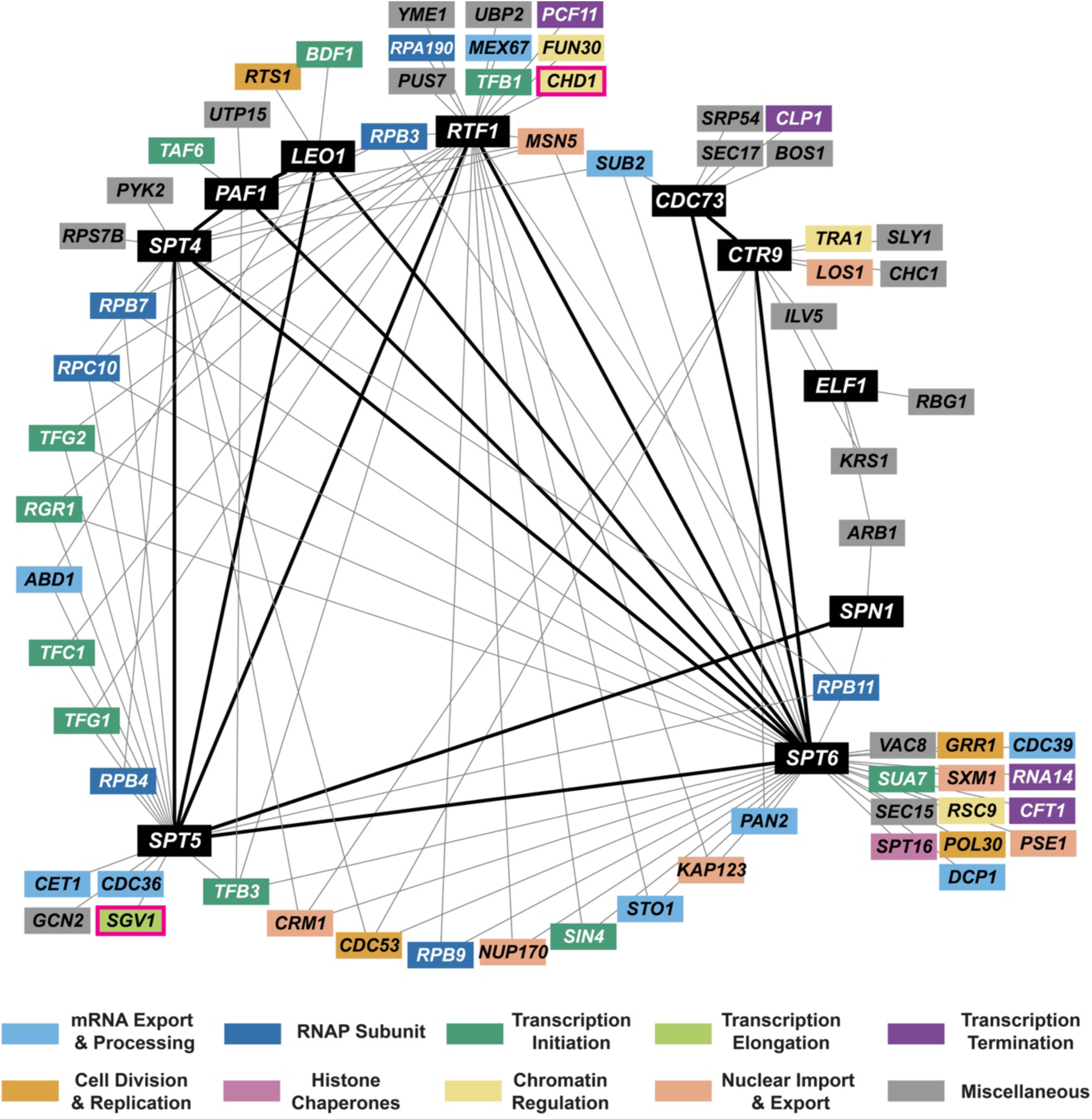
ERC analysis identifies known and putative functional interactors of TEFs. Each node in the network represents a gene, and an edge between two nodes indicates that the ERC between the two genes has a Z-score greater than 3.5. Black nodes represent TEFs, and black edges represent connections among TEFs. ERC values between genes in this network range from 7.6 to 16.0. Color of nodes represents the function attributed to the gene. Miscellaneous functional categories include metabolism (*PYK2, ILV5, PUS7*), protein homeostasis (*YME1, UBP2*), protein trafficking (*VAC8, CHC1, SRP54*), ribosome subunits and biogenesis (*RPS7B, ARB1*), rRNA processing (*UTP15*), translation (*KRS1, RBG1, GCN2*) and vesicle trafficking (*SEC15, SLY1, SEC17, BOS1*). Boxed in pink are nodes of interest that are discussed in the main text.

As predicted, several known functional interactors of TEFs share a high ERC with them. Here, we highlight two examples (nodes highlighted by pink outlines in **Fig. 5**). First, the chromatin remodeler *CHD1* has a high ERC with *RTF1* (Z-score*_RTF1-CHD1_* = 3.88). As noted above, the LLALA box in Rtf1 directly interacts with the CHCT domain of Chd1 and is required for proper localization of Chd1 on gene bodies (43, 57, 58). Second, *SGV1*/*BUR1* has a high ERC with *SPT5* (Z-score*_SPT5-SGV1_* = 3.56). Bur1 phosphorylates the C-terminal repeat region of Spt5, and this phosphorylation is required for the proper recruitment of Paf1C onto gene bodies (59–62).

Using the Z-score cut-off, we also note that compared to other TEFs, a greater number of genes have a high ERC with *SPT6* and *RTF1* (**Fig. S12B**). Additionally, a high proportion of genes co-varying with *SPT6, RTF1,* and *CDC73* are unique to these TEFs (**Fig. S12B**), suggestive of individualized functions not generally shared between TEFs. These include, but are not restricted to, genes encoding several transcription termination factors (Pcf11, Clp1, Rna14, and Cft1; purple nodes in **Fig. 5**). Overall, this analysis highlights both known and underappreciated functions of TEFs and identifies putative functional interactors that warrant further experimental characterization.

## Discussion

### TEFs are broadly conserved across eukaryotes with notable exceptions

This study provides a comprehensive analysis of core TEF orthologs in 304 species distributed across the Tree of Life to understand variations and conserved features of eukaryotic transcription regulation (**Fig. 6**). Our findings demonstrate the deep evolutionary origins of TEFs, reveal key architectural and regulatory features that have been maintained for over a billion years, and uncover lineage-specific innovations. As expected, we find Spt5 in all domains of life and Spt4 and Elf1 orthologs in both Archaea and Eukaryota. All subunits of Paf1C and Spn1 appear to be eukaryote-specific. Their presence in early branching eukaryotic lineages indicates establishment prior to LECA. Paf1C subunits were less consistently detected by HMM search, particularly in discobid, metamonad, and alveolate proteomes.

**Figure 6.**
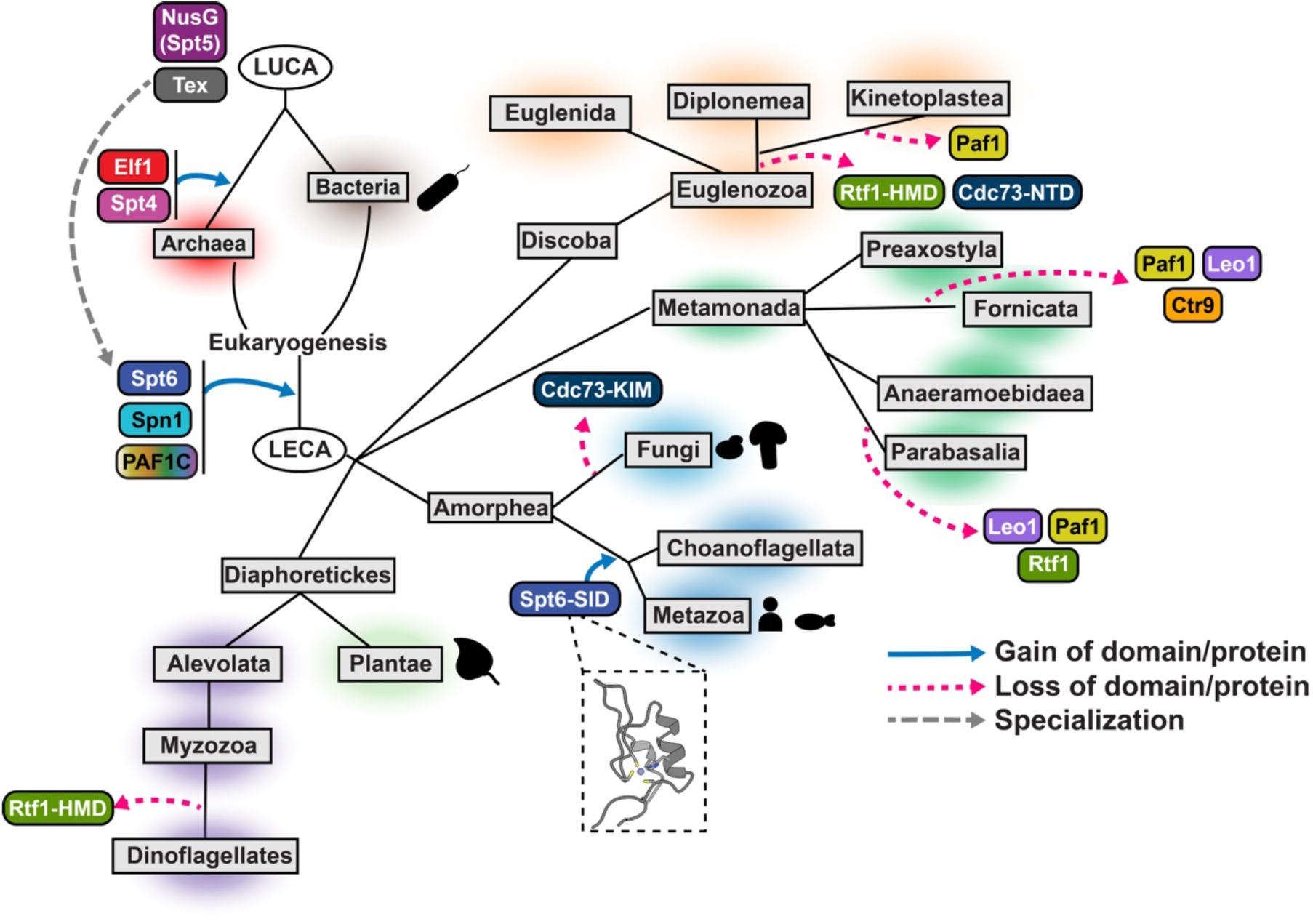
Model of TEF domain evolution. Proposed model of acquisition and loss of TEF domains across clades in the Tree of Life. Spt5 is the only known universal elongation factor. Spt4 and Elf1 are archaeal innovations. Spt6, Spn1, and the Paf1C are eukaryotic-elongation factors. Though broadly conserved, we were unable to detect specific Paf1C domains in some eukaryotic clades. Spt6 homologs in choanoflagellates and metazoans have gained a zinc finger domain we termed the SID.

The reasons for the apparent dispensability of Paf1C domains within some clades remain unclear. However, in the case of the Rtf1 HMD, this variability may be linked to the key role of the HMD in stimulating H2B monoubiquitylation (H2Bub) (58, 63, 64). We speculate that altered chromatin biology and transcriptional regulation in these clades may relax the evolutionary constraints coordinating H2Bub with transcription elongation, leading to the divergence or loss of the Rtf1 HMD. Furthermore, in clades where HMD is not detectable, alternative mechanisms may have evolved to compensate for the loss of HMD-dependent functions. For example, in Opisthokonts, H2Bub is required for co-transcriptional H3K79 methylation by the Dot1 methyltransferase (65–67). In kinetoplastids, a sub-clade of Discoba in which the HMD and H2Bub are not detected, Dot1 orthologs have been reported to operate independently of H2Bub (68).

### A repertoire of conserved TEF features

Mapping conservation scores onto protein structures yields functional maps of TEFs that suggest testable mechanisms. While we demonstrate that this analysis can sensitively highlight known protein-protein interfaces (**Figs. 3B** and **3C**), we find no such conservation hotspots along the length of the Ctr9 trestle. These observations suggest this helix is primarily an architectural feature or acts to sterically occlude portions of RNAPII from other factors, though further work is needed to test these functions. Comparative conservation score analyses additionally identify concordantly and discordantly conserved regions of fungal and metazoan TEFs. We also identify several poorly understood structural features of TEFs, including regions along Paf1, Ctr9, Cdc73, Rtf1, and Spt6 (**Fig. 4**). As a resource for future studies, ChimeraX structural visualization session files highlighting conserved regions in these proteins have been provided (**Supplementary File**) (69). This evolutionary framework should accelerate hypothesis-driven mutagenesis experiments, unify findings across model systems, and guide the discovery of lineage-specific mechanisms of transcription elongation regulation.

### The under-characterized features of the transcription elongation complex

TEFs have been studied for decades, yet our results indicate that there remains much to explore. Amongst the most conspicuous and uncharacterized regions in metazoan TEFs are the Spt6 SID, Cdc73 NTD, and the N-terminal extension from the Rtf1 LLALA box. Combining our insights with analyses enabled by powerful methodologies such as targeted AlphaFold multimer screens (70), site-specific crosslinking (71), and classical biochemical and genetic approaches will be crucial to clarify the specific biological functions of TEFs.

ERC analyses of yeast species further suggest underexplored connections in the evolutionary patterns shared amongst TEFs (**Fig. 5**). Intriguingly, evolutionary rates of *SPT6*, *RTF1*, and *SPT5* co-vary with a high number of genes, and many genes exhibit evolutionary rates uniquely co-varying with *RTF1*, *SPT6*, or *CDC73* (**Fig. S12B**). We propose two non-exclusive models to explain our observations. First, these TEFs might serve as multifunctional hubs in the elongation complex, bridging the physical or functional interactions of diverse proteins with the transcription elongation machinery. This is consistent with several studies implicating the requirement of Spt6, Rtf1, and Cdc73 for proper transcription termination (2, 72–75). Second, these TEFs might play an important role in other cellular processes not directly related to transcription elongation. In support of this model, *SPT6* has a high ERC with *POL30* (Z-score*_SPT6-POL30_* = 3.66), which encodes the DNA polymerase processivity factor PCNA (76). Consistent with this observation, Spt6 is required for DNA replication and genome stability in *S. cerevisiae* (2, 74, 77). We speculate that high ERC pairs such as *RTF1* and the *FUN30,* which encodes a chromatin remodeler, or *LEO1* and the *RTS1,* which encodes a regulatory subunit of PP2A phosphatase, represent candidate functional interactors with the elongation machinery.

While informative, our approach is limited in its ability to parse function from rapidly evolving disordered regions, such as those involved in histone interactions or phase separation (29, 78, 79). Additionally, several reported interactions, including those between Spt6 and Spn1 (80–82) and between Cdc73 and Spt6 (83, 84), are unresolved in the 9EH2 PDB structure and as such remain unexamined in this study. Nevertheless, this work provides a rich, evolution-guided roadmap for future mechanistic studies to understand conserved and divergent principles of transcription regulation.

## Materials and Methods

### Data retrieval and domain annotations

Domain annotations were manually curated from published crystal or cryo-EM structures of *H. sapiens* or *S. cerevisiae* proteins or, when these were not available, from structures predicted by AlphaFold. Predicted protein structures were downloaded from the AlphaFold Protein Structure Database (December 12, 2023) (85, 86). Domain boundaries and the sources of their annotations are provided in Zenodo repository (see **Data and code accessibility**).

Prokaryotic proteome sequences were retrieved from the Genome Taxonomy Database (GTDB) (release 214, April 21, 2023) (87–90). For the combined EukProt/GTDB proteome search, Archaeal (n = 37) and Bacterial (n = 40) proteome sequences representing a range of prokaryotic clades were manually subset from GTDB to be included in the search. Only Prokaryotic proteomes annotated as “GTDB species representatives” and “NCBI type material” were considered. Eukaryotic proteomes (n = 227) were retrieved from the EukProt Database (Version 3, November 22, 2022) (91). Eukaryotic species selected included the full “Comparative Set”, which had been curated to represent maximum breadth and proteome completeness (91).

For a deeper search within Asgardarchaeota, 218 proteome sequences were subset from the same GTDB release. Additional RefSeq annotated Fungal (n=128) and Metazoan (n=706) proteomes (assembly level – chromosome and complete) were downloaded from NCBI RefSeq (January 9, 2025) (92). Species lists of each proteome set are provided in Zenodo repository (see **Data and code accessibility**).

### Domain-forward search for elongation factor homologs

Searches for orthologs of human TEFs were performed using BLAST+ v2.13.0 (93) against proteomes from GTDB and EukProt (-evalue 10). The hits with the lowest e-value for each species were then aligned using MUSCLE (v5.1) (94) (-super5). Next, we individually searched for the domains of each protein through the proteome database. To do this, we first trimmed the relevant MSA to the boundaries of the domain of interest as annotated in the *S. cerevisiae* and *H. sapiens* orthologs of interest using a custom script. Entries from species that showed no protein sequence over the portion of the MSA containing that domain were purged from the domain-trimmed MSA using seqkit (v0.16.0) (95, 96) (seqkit grep --invert-match -r -p “-+$”). The resultant alignment was used to build an HMM using hmmbuild from the hmmer package (v3.2.1) (97). The combined GTDB and EukProt database was probed for matches to each domain HMM with the hmmsearch command (minimum e-value = 10^-3^) using the “wrap_hmmscan.pl” script produced by Dan Richter (available at https://github.com/MBL-Physiology-Bioinformatics/2021-Bioinformatics-Tutorial-Materials/tree/master/phylogenetics).

The top hit for each species was collected and realigned to the original HMM using hmmer hmmalign. Using a custom script to filter this new alignment, sequences with information-poor poly(K)/poly(Q) stretches >10 residues long or insertions that were not shared by 5% or more of other species were omitted. The remaining sequences were once again realigned using hmmalign, and a new HMM was built. This HMM was used to search the proteome database as before, this time using a minimum e-value of 10^-4^ and retaining the top 3 hits of potential homologs.

For each protein, the domain-specific HMM search hits and original BLAST search hits were then compiled into one FASTA file and duplicate sequences removed with seqkit rmdup (--by-name). To validate that the putative homolog hits were most closely related to the proteins of interest, we performed a reciprocal BLAST against *S. cerevisiae*, *H. sapiens*, and *A. thaliana* proteomes. Hits with the greatest similarity (lowest e-value) to known annotations of the protein of interest were considered “putatively valid”. Hits that showed similarity to the protein of interest but a greater similarity to a different protein in the model species were considered “potentially valid”. If hits showed greater similarity to an unrelated protein known to bear a similar structure (*e.g*. TPR containing protein Cyc8 shares similarities with Ctr9), they were blacklisted and filtered out. Hits that showed no significant similarity to the protein of interest were considered dubious. Potentially valid and dubious hits were excluded if they showed no significant similarity to the protein of interest by either NCBI BLAST or PFAM scans of the sequence (23). The remaining sequences were aligned with the addition of outgroup sequences using MUSCLE (v5.1) (-super5) and trimmed using ClipKIT (v1.4.1) (98) (-m kpi-gappy). Phylogenetic trees were constructed with both FastTree (v2.1.10) (99) (default settings) and iqtree2 (v2.2.2.7) (100, 101) (-m MFP -B 1000 --mem 250G -T AUTO). Outgroups and false-positive hits were manually removed to provide a final list of probable homologs. Manual removal was performed by inspection of domain architecture (predicted using PFAM scans) and gene trees to yield a final set of filtered orthologs. Since the S1 domain of Spt6 is common amongst RNA-binding proteins, HMM searches for the S1 domain were poorly selective for Spt6 and were thus excluded from the analysis. Blacklisted proteins and outgroup proteins are provided in Zenodo repository (see **Data and code accessibility**).

To search for putative TEF orthologs in Archaea, we performed scans with each finalized TEF domain HMM against GTDB Asgardarchaeota proteomes (minimum e-value = 10^-3^) and filtered by reciprocal blast as above, retaining only hits with greatest similarity to known reference TEFs. For Metazoan or Fungi targeted searches, a BLAST search was performed using human and yeast orthologs against their respective databases (-max_target_seqs 50000) and only the hit with the lowest e-value for each species was retained, followed by alignment with MUSCLE (v5.1) (-super5). Filtered orthologs identified in the Eukprot/GTDB database search were aligned with MUSCLE (v5.1) (-super5) and a custom script was used to identify the mapping of analogous residues between *H. sapiens* and *S. cerevisiae* TEF reference sequences.

Details of additional analyses and conservation score calculations are described in the supplementary methods.

## Acknowledgments

We are grateful to Tera Levin and Ed Culbertson for their advice on the HMM-based search for TEF orthologs and helpful discussions. We thank Sarah Hainer, Craig Kaplan, Miler Lee, members of their groups, and all members of the Arndt lab for insightful discussions. We are grateful to Fred Winston, Craig Kaplan, and members of the Arndt lab for their critical reading of the manuscript. This research was supported in part by the University of Pittsburgh Center for Research Computing and Data, RRID:SCR_022735, through the resources provided. Specifically, this work used the HTC cluster, which is supported by NIH award number S10OD028483, and the H2P cluster, which is supported by NSF award number OAC-2117681. This work was supported by an Andrew Mellon Predoctoral fellowship to A.G., a Provost’s Dissertation Year Fellowship for Historically Underrepresented Doctoral Students to A.M.F., NIH grant R01 HG009299 to N.C, and NIH grant R35 GM141964 to K.M.A.

## Author Contributions

A.M.F, A.G., N.C., and K.M.A. designed research; A.M.F and A.G. performed research; A.M.F, A.G., N.C., and K.M.A. analyzed data; A.M.F and A.G. contributed new analytical tools; and A.M.F, A.G., and K.M.A. wrote the paper.

## References

1. A. M. Francette, K. M. Arndt, Multiple direct and indirect roles of the Paf1 complex in transcription elongation, splicing, and histone modifications. Cell Rep 43, 114730 (2024).

2. C. L. W. Miller, J. L. Warner, F. Winston, Insights into Spt6: a histone chaperone that functions in transcription, DNA replication, and genome stability. Trends Genet 39, 858–872 (2023).

3. T. M. Decker, Mechanisms of Transcription Elongation Factor DSIF (Spt4-Spt5). J Mol Biol 433, 166657 (2021).

4. L. Farnung, Chromatin Transcription Elongation - A Structural Perspective. J Mol Biol 437, 168845 (2025).

5. A. Zheenbekova et al., IWS1 positions downstream DNA to globally stimulate Pol II elongation. Nat Commun 16, 7747 (2025).

6. D. Syau et al., Structure and function of IWS1 in transcription elongation. bioRxiv 10.1101/2025.08.28.672863 (2025).

7. F. Burki, A. J. Roger, M. W. Brown, A. G. B. Simpson, The New Tree of Eukaryotes. Trends Ecol Evol 35, 43–55 (2020).

8. G. Lax et al., Hemimastigophora is a novel supra-kingdom-level lineage of eukaryotes. Nature 564, 410–414 (2018).

9. G. Torruella, L. J. Galindo, D. Moreira, P. Lopez-Garcia, Phylogenomics of neglected flagellated protists supports a revised eukaryotic tree of life. Curr Biol 35, 198–207 e194 (2025).

10. T. Fouqueau et al., The cutting edge of archaeal transcription. Emerg Top Life Sci 2, 517–533 (2018).

11. C. P. Ponting, Novel domains and orthologues of eukaryotic transcription elongation factors. Nucleic Acids Res 30, 3643–3652 (2002).

12. T. Wada et al., DSIF, a novel transcription elongation factor that regulates RNA polymerase II processivity, is composed of human Spt4 and Spt5 homologs. Genes Dev 12, 343–356 (1998).

13. M. Guo et al., Core structure of the yeast spt4-spt5 complex: a conserved module for regulation of transcription elongation. Structure 16, 1649–1658 (2008).

14. F. Blombach, T. Fouqueau, D. Matelska, K. Smollett, F. Werner, Promoter-proximal elongation regulates transcription in archaea. Nat Commun 12, 5524 (2021).

15. C. D. Kaplan, J. R. Morris, C. Wu, F. Winston, Spt5 and spt6 are associated with active transcription and have characteristics of general elongation factors in D. melanogaster. Genes Dev 14, 2623–2634 (2000).

16. D. Close et al., Crystal structures of the S. cerevisiae Spt6 core and C-terminal tandem SH2 domain. J Mol Biol 408, 697–713 (2011).

17. S. J. Johnson et al., Crystal structure and RNA binding of the Tex protein from Pseudomonas aeruginosa. J Mol Biol 377, 1460–1473 (2008).

18. A. G. Chivu et al., Evolution of promoter-proximal pausing enabled a new layer of transcription control. Nat Struct Mol Biol 10.1038/s41594-025-01718-y (2025).

19. B. A. Ouna et al., Depletion of Trypanosome CTR9 Leads to Gene Expression Defects. PLOS ONE 7, e34256 (2012).

20. P. Mitra, A. S. Deshmukh, S. Banerjee, C. Khandavalli, C. Choudhury, A functionally divergent transcription elongation factor 1-like protein in Toxoplasma gondii. FEBS Letters 596, 112–127 (2022).

21. A. Srivastava, N. Badjatia, J. H. Lee, B. Hao, A. Gunzl, An RNA polymerase II-associated TFIIF-like complex is indispensable for SL RNA gene transcription in Trypanosoma brucei. Nucleic Acids Res 46, 1695–1709 (2018).

22. S. Chanarat, Transcription machinery of the minimalist: comparative genomic analysis provides insights into the (de)regulated transcription mechanism of microsporidia - fungal-relative parasites. Transcription 14, 1–17 (2023).

23. J. Mistry et al., Pfam: The protein families database in 2021. Nucleic Acids Res 49, D412–D419 (2021).

24. J. P. Daniels, S. Kelly, B. Wickstead, K. Gull, Identification of a crenarchaeal orthologue of Elf1: implications for chromatin and transcription in Archaea. Biol Direct 4, 24 (2009).

25. J. Vosseberg et al., The emerging view on the origin and early evolution of eukaryotic cells. Nature 633, 295–305 (2024).

26. J. W. Markert, J. H. Soffers, L. Farnung, Structural basis of H3K36 trimethylation by SETD2 during chromatin transcription. Science 387, 528–533 (2025).

27. I. Friedberg, H. Margalit, Persistently conserved positions in structurally similar, sequence dissimilar proteins: roles in preserving protein fold and function. Protein Sci 11, 350–360 (2002).

28. S. M. Vos, L. Farnung, A. Linden, H. Urlaub, P. Cramer, Structure of complete Pol II-DSIF-PAF-SPT6 transcription complex reveals RTF1 allosteric activation. Nat Struct Mol Biol 27, 668–677 (2020).

29. H. Lyons et al., Functional partitioning of transcriptional regulators by patterned charge blocks. Cell 186, 327–345 e328 (2023).

30. P. W. Chiang et al., Isolation and characterization of the human and mouse homologues (SUPT4H and Supt4h) of the yeast SPT4 gene. Genomics 34, 368–375 (1996).

31. H. Liu et al., A taxon-rich and genome-scale phylogeny of Opisthokonta. PLoS Biol 22, e3002794 (2024).

32. N. A. O’Leary et al., Reference sequence (RefSeq) database at NCBI: current status, taxonomic expansion, and functional annotation. Nucleic Acids Res 44, D733–745 (2016).

33. J. L. Walshe et al., Molecular mechanism of co-transcriptional H3K36 methylation by SETD2. Nat Commun 16, 9565 (2025).

34. Y. Xie et al., Paf1 and Ctr9 subcomplex formation is essential for Paf1 complex assembly and functional regulation. Nat Commun 9, 3795 (2018).

35. S. M. Vos et al., Structure of activated transcription complex Pol II-DSIF-PAF-SPT6. Nature 560, 607–612 (2018).

36. P. Deng et al., Transcriptional elongation factor Paf1 core complex adopts a spirally wrapped solenoidal topology. Proc Natl Acad Sci U S A 115, 9998–10003 (2018).

37. W. Sun et al., Crystal structure of the N-terminal domain of human CDC73 and its implications for the hyperparathyroidism-jaw tumor (HPT-JT) syndrome. Sci Rep 7, 15638 (2017).

38. B. Zhu et al., The human PAF complex coordinates transcription with events downstream of RNA synthesis. Genes Dev 19, 1668–1673 (2005).

39. D. Lopez Martinez et al., PAF1C-mediated activation of CDK12/13 kinase activity is critical for CTD phosphorylation and transcript elongation. Mol Cell 85, 1952–1967 e1958 (2025).

40. M. A. Hahn, D. J. Marsh, Nucleolar localization of parafibromin is mediated by three nucleolar localization signals. FEBS Lett 581, 5070–5074 (2007).

41. M. A. Hahn, D. J. Marsh, Identification of a functional bipartite nuclear localization signal in the tumor suppressor parafibromin. Oncogene 24, 6241–6248 (2005).

42. Y. J. Yang, J. W. Han, H. D. Youn, E. J. Cho, The tumor suppressor, parafibromin, mediates histone H3 K9 methylation for cyclin D1 repression. Nucleic Acids Res 38, 382–390 (2010).

43. S. A. Tripplehorn et al., A direct interaction between the Chd1 CHCT domain and Rtf1 controls Chd1 distribution and nucleosome positioning on active genes. Nucleic Acids Res 53 (2025).

44. M. Bycroft, T. J. Hubbard, M. Proctor, S. M. Freund, A. G. Murzin, The solution structure of the S1 RNA binding domain: a member of an ancient nucleic acid-binding fold. Cell 88, 235–242 (1997).

45. B. F. Lang, C. O’Kelly, T. Nerad, M. W. Gray, G. Burger, The closest unicellular relatives of animals. Curr Biol 12, 1773–1778 (2002).

46. F. Chen et al., Crystal Structure of the Core Module of the Yeast Paf1 Complex. J Mol Biol 434, 167369 (2022).

47. Y. Qin et al., Structural Basis of the Transcriptional Elongation Factor Paf1 Core Complex from Saccharomyces eubayanus. International Journal of Molecular Sciences 24, 8730 (2023).

48. H. Ehara, T. Kujirai, M. Shirouzu, H. Kurumizaka, S. I. Sekine, Structural basis of nucleosome disassembly and reassembly by RNAPII elongation complex with FACT. Science 377, eabp9466 (2022).

49. K. Adelman et al., *Drosophila* Paf1 modulates chromatin structure at actively transcribed genes. Mol Cell Biol 26, 250–260 (2006).

50. J. Kim, M. Guermah, R. G. Roeder, The human PAF1 complex acts in chromatin transcription elongation both independently and cooperatively with SII/TFIIS. Cell 140, 491–503 (2010).

51. A. D. Langenbacher et al., The PAF1 complex differentially regulates cardiomyocyte specification. Dev Biol 353, 19–28 (2011).

52. O. Rozenblatt-Rosen et al., The parafibromin tumor suppressor protein is part of a human Paf1 complex. Mol Cell Biol 25, 612–620 (2005).

53. Q. F. Cao et al., Characterization of the human transcription elongation factor Rtf1: Evidence for nonoverlapping functions of Rtf1 and the Paf1 complex. Mol Cell Biol 35, 3459–3470 (2015).

54. N. L. Clark, E. Alani, C. F. Aquadro, Evolutionary rate covariation reveals shared functionality and coexpression of genes. Genome Res 22, 714–720 (2012).

55. G. J. Brunette, M. A. Jamalruddin, R. A. Baldock, N. L. Clark, K. A. Bernstein, Evolution-based screening enables genome-wide prioritization and discovery of DNA repair genes. Proc Natl Acad Sci U S A 116, 19593–19599 (2019).

56. J. H. Little et al., ERC2.0 evolutionary rate covariation update improves inference of functional interactions across large phylogenies. Genome Res 10.1101/gr.280586.125 (2025).

57. R. Simic et al., Chromatin remodeling protein Chd1 interacts with transcription elongation factors and localizes to transcribed genes. EMBO J 22, 1846–1856 (2003).

58. M. H. Warner, K. L. Roinick, K. M. Arndt, Rtf1 is a multifunctional component of the Paf1 complex that regulates gene expression by directing cotranscriptional histone modification. Mol Cell Biol 27, 6103–6115 (2007).

59. K. Zhou, W. H. Kuo, J. Fillingham, J. F. Greenblatt, Control of transcriptional elongation and cotranscriptional histone modification by the yeast BUR kinase substrate Spt5. Proc Natl Acad Sci U S A 106, 6956–6961 (2009).

60. Y. Liu et al., Phosphorylation of the transcription elongation factor Spt5 by yeast Bur1 kinase stimulates recruitment of the PAF complex. Mol Cell Biol 29, 4852–4863 (2009).

61. M. K. Mayekar, R. G. Gardner, K. M. Arndt, The recruitment of the Saccharomyces cerevisiae Paf1 complex to active genes requires a domain of Rtf1 that directly interacts with the Spt4-Spt5 complex. Mol Cell Biol 33, 3259–3273 (2013).

62. S. Namjilsuren, K. M. Arndt, The Glc7/PP1 phosphatase triggers Paf1C dissociation from RNA polymerase II to enable transcription termination. bioRxiv 10.1101/2025.09.03.674035, 2025.2009.2003.674035 (2025).

63. S. B. Van Oss et al., The Histone Modification Domain of Paf1 Complex Subunit Rtf1 Directly Stimulates H2B Ubiquitylation through an Interaction with Rad6. Mol Cell 64, 815–825 (2016).

64. A. S. Piro, M. K. Mayekar, M. H. Warner, C. P. Davis, K. M. Arndt, Small region of Rtf1 protein can substitute for complete Paf1 complex in facilitating global histone H2B ubiquitylation in yeast. Proc Natl Acad Sci U S A 109, 10837–10842 (2012).

65. M. D. Shahbazian, K. Zhang, M. Grunstein, Histone H2B ubiquitylation controls processive methylation but not monomethylation by Dot1 and Set1. Mol Cell 19, 271–277 (2005).

66. H. H. Ng, R. M. Xu, Y. Zhang, K. Struhl, Ubiquitination of histone H2B by Rad6 is required for efficient Dot1-mediated methylation of histone H3 lysine 79. J Biol Chem 277, 34655–34657 (2002).

67. R. K. McGinty, J. Kim, C. Chatterjee, R. G. Roeder, T. W. Muir, Chemically ubiquitylated histone H2B stimulates hDot1L-mediated intranucleosomal methylation. Nature 453, 812–816 (2008).

68. V. S. Frisbie et al., Two DOT1 enzymes cooperatively mediate efficient ubiquitin-independent histone H3 lysine 76 tri-methylation in kinetoplastids. Nature Communications 15, 2467 (2024).

69. E. C. Meng et al., UCSF ChimeraX: Tools for structure building and analysis. Protein Sci 32, e4792 (2023).

70. D. Molodenskiy et al., AlphaPulldown2-a general pipeline for high-throughput structural modeling. Bioinformatics 41 (2025).

71. J. W. Chin, A. B. Martin, D. S. King, L. Wang, P. G. Schultz, Addition of a photocrosslinking amino acid to the genetic code of Escherichiacoli. Proc Natl Acad Sci U S A 99, 11020–11024 (2002).

72. O. Rozenblatt-Rosen et al., The tumor suppressor Cdc73 functionally associates with CPSF and CstF 3’ mRNA processing factors. Proc Natl Acad Sci U S A 106, 755–760 (2009).

73. B. N. Tomson, C. P. Davis, M. H. Warner, K. M. Arndt, Identification of a role for histone H2B ubiquitylation in noncoding RNA 3’-end formation through mutational analysis of Rtf1 in Saccharomyces cerevisiae. Genetics 188, 273–289 (2011).

74. A. Narain et al., Targeted protein degradation reveals a direct role of SPT6 in RNAPII elongation and termination. Mol Cell 81, 3110–3127 e3114 (2021).

75. Y. Yang et al., PAF Complex Plays Novel Subunit-Specific Roles in Alternative Cleavage and Polyadenylation. PLoS Genet 12, e1005794 (2016).

76. L. M. Dieckman, B. D. Freudenthal, M. T. Washington, PCNA structure and function: insights from structures of PCNA complexes and post-translationally modified PCNA. Subcell Biochem 62, 281–299 (2012).

77. R. Franklin et al., Histone chaperones coupled to DNA replication and transcription control divergent chromatin elements to maintain cell fate. Genes Dev 39, 652–675 (2025).

78. C. Evrin et al., Spt5 histone binding activity preserves chromatin during transcription by RNA polymerase II. EMBO J 41, e109783 (2022).

79. A. Kasiliauskaite et al., Cooperation between intrinsically disordered and ordered regions of Spt6 regulates nucleosome and Pol II CTD binding, and nucleosome assembly. Nucleic Acids Res 50, 5961–5973 (2022).

80. N. J. Krogan et al., RNA polymerase II elongation factors of Saccharomyces cerevisiae: a targeted proteomics approach. Mol Cell Biol 22, 6979–6992 (2002).

81. S. M. McDonald, D. Close, H. Xin, T. Formosa, C. P. Hill, Structure and biological importance of the Spn1-Spt6 interaction, and its regulatory role in nucleosome binding. Mol Cell 40, 725–735 (2010).

82. M. L. Diebold et al., The structure of an Iws1/Spt6 complex reveals an interaction domain conserved in TFIIS, Elongin A and Med26. EMBO J 29, 3979–3991 (2010).

83. M. A. Ellison et al., Spt6 directly interacts with Cdc73 and is required for Paf1 complex occupancy at active genes in Saccharomyces cerevisiae. Nucleic Acids Res 51, 4814–4830 (2023).

84. L. Zhang et al., Structural basis of RECQL5-induced RNA polymerase II transcription braking and subsequent reactivation. Nat Struct Mol Biol 10.1038/s41594-025-01586-6 (2025).

85. J. Jumper et al., Highly accurate protein structure prediction with AlphaFold. Nature 596, 583–589 (2021).

86. M. Varadi et al., AlphaFold Protein Structure Database in 2024: providing structure coverage for over 214 million protein sequences. Nucleic Acids Res 52, D368–D375 (2024).

87. D. H. Parks et al., GTDB: an ongoing census of bacterial and archaeal diversity through a phylogenetically consistent, rank normalized and complete genome-based taxonomy. Nucleic Acids Res 50, D785–D794 (2022).

88. C. Rinke et al., A standardized archaeal taxonomy for the Genome Taxonomy Database. Nat Microbiol 6, 946–959 (2021).

89. D. H. Parks et al., A complete domain-to-species taxonomy for Bacteria and Archaea. Nat Biotechnol 38, 1079–1086 (2020).

90. D. H. Parks et al., A standardized bacterial taxonomy based on genome phylogeny substantially revises the tree of life. Nat Biotechnol 36, 996–1004 (2018).

91. D. Richter, et al., EukProt: A database of genome-scale predicted proteins across the diversity of eukaryotes. Peer Community Journal 2 (2022).

92. T. Goldfarb et al., NCBI RefSeq: reference sequence standards through 25 years of curation and annotation. Nucleic Acids Res 53, D243–D257 (2025).

93. C. Camacho et al., BLAST+: architecture and applications. BMC Bioinformatics 10, 421 (2009).

94. R. C. Edgar, MUSCLE: multiple sequence alignment with high accuracy and high throughput. Nucleic Acids Res 32, 1792–1797 (2004).

95. W. Shen, S. Le, Y. Li, F. Hu, SeqKit: A Cross-Platform and Ultrafast Toolkit for FASTA/Q File Manipulation. PLoS One 11, e0163962 (2016).

96. W. Shen, B. Sipos, L. Zhao, SeqKit2: A Swiss army knife for sequence and alignment processing. Imeta 3, e191 (2024).

97. S. C. Potter et al., HMMER web server: 2018 update. Nucleic Acids Res 46, W200–W204 (2018).

98. J. L. Steenwyk, T. J. Buida, 3rd, Y. Li, X. X. Shen, A. Rokas, ClipKIT: A multiple sequence alignment trimming software for accurate phylogenomic inference. PLoS Biol 18, e3001007 (2020).

99. M. N. Price, P. S. Dehal, A. P. Arkin, FastTree 2--approximately maximum-likelihood trees for large alignments. PLoS One 5, e9490 (2010).

100. B. Q. Minh et al., IQ-TREE 2: New Models and Efficient Methods for Phylogenetic Inference in the Genomic Era. Mol Biol Evol 37, 1530–1534 (2020).

101. D. T. Hoang, O. Chernomor, A. von Haeseler, B. Q. Minh, L. S. Vinh, UFBoot2: Improving the Ultrafast Bootstrap Approximation. Mol Biol Evol 35, 518–522 (2018).

