## Supplemental Material for "Evolutionary conservation and innovations of RNA polymerase II transcription elongation factors"

### Supplementary Methods

#### Alternative search strategies for TEF homologs

To search for TEF homologs in specific sub-clades, a three-step approach was used. In clades where a given domain was not detected in any species by the HMM searches, FoldSeek was used to detect domains using structural similarity. FoldSeek searches were performed on the web server (1). When available, existing crystal structures of domains were used as inputs for the search - Rtf1-HMD (PDB ID 5E8B) (2), Cdc73-NTD (PDB ID 5YDE) (3), Ctr9-TPR motifs (PDB ID 6AF0) (4) and Rtf1-Plus3 (PBD ID 4L1P) (5). In other cases, AlphaFold-predicted structures trimmed to the domain boundaries were used as inputs. Searches were performed against the AlphaFold/Uniprot50 v4 database by filtering for the clade indicated in the Zenodo repository (see **Supplementary Table 1** and **Data and code accessibility**). The remaining parameters were set to default. In cases where FoldSeek was unable to detect the domain of interest, AlphaFold3 web server (6) was used to predict the structures of candidate orthologs. Predicted structures were then manually inspected for the presence of the domain of interest. In cases where no protein hit corresponding to the protein of interest was identified by the HMM search, this was not carried out. When FoldSeek and AlphaFold3 were unable to detect the domain of interest, PSI-BLAST was performed. PSI-BLAST was performed on the BLAST web server (7) and was carried out using default parameters with the following exceptions: the Database was set to ClusteredNR, Organism was set to taxa indicated in **Supplementary Table 1**. Max target sequences was set to 20000, and Expect threshold was set to 1. To maximize our chances of finding these domains using PSI-BLAST in the mentioned taxa, hits from closely related clades were used as templates.

#### Relative evolutionary rate calculations and structural representations

Each final set of homologs (EukProt + GTDB, refseq Metazoan, and refseq Fungi) was aligned using MAFFT (v7.471) (--auto –anysymbol), then MSAs were trimmed using ClipKIT (v1.4.1) (-m kpi-gappy). The trimmed MSAs were used to build trees using iqtree2 (-m MFP -B 1000 --mem 250G -T AUTO). Separately, copies of the original MSA were processed such that columns not represented in the reference of interest (*H. sapiens* or *S. cerevisiae*) were discarded using a custom script. MEGA11 (8) was used to calculate relative evolutionary rates (RERs). To do so, iqtree2 TEF gene trees and MAFFT MSAs (condensed to columns represented in the *H. sapiens* or *S. cerevisiae* TEF reference as indicated in text) were used as input for the “Estimate Rate at Each Site (ML)” analysis (- Statistical Method “Maximum Likelihood” - Substitutions Type “Amino Acid” - Model/Method “JTT” - Rates among sites “Gamma Distributed” - No. of Discrete Gamma Categories “5”). All sites were used for the calculation, and no branch swap filter was applied. RERs are calculated such that RER = 0 is invariant, RER = 1 is an average rate of evolution, and RER > 1 is a greater than average rate of evolution. Conservation scores were calculated as -log10(RER + 0.1). Final modified and unmodified MSAs, gene trees, RERs, and conservation scores are provided in Zenodo repository.

To map RERs onto molecular structures the ß-factors of the AlphaFold predictions of the relevant monomers were overwritten using a custom script. We have provided the associated sessions in the open-source ChimeraX viewer (v1.9) (9) as supplementary files in Zenodo repository (see **Data and code accessibility**). Per-residue surface-accessible surface area (SASA) and surface lipophilicity maps were calculated in ChimeraX. SASA measurements were based on each TEF’s monomeric AlphaFold predicted structure and residues with SASA > 50 $Å^2$ were considered “solvent-exposed”.

**AlphaFold3 structure predictions**

AlphaFold3 (6) was used to predict the structure of putative TEF homologs (**Fig. 2B-D**, **Fig. S7A**, and **Supplementary Table 1**) and the *H. sapiens* Spt6 SID (**Fig. S10A-B**) and to model the putative interaction interface between Rtf1 and Ctr9 (**Fig. S11C-G**). *H. sapiens* Spt6 S1 domain, along with a $Zn^{2+}$ ion, was used as input for the prediction. *H. sapiens* and *S. cerevisiae* Rtf1-Ctr9 multimer predictions were carried out using 10 different seeds (10, 15, 500, 43, 58, 5697, 4327, 798, 111, 287), with each seed yielding 5 models. Structures were visualized in ChimeraX.

**S1 insert size distribution analysis**

A custom script was used to calculate the size of SID-like insertions within known Spt6 orthologs. Briefly, the MUSCLE-aligned MSA of top-scoring Spt6 orthologs from the Eukprot/GTDB search was loaded into R using the msa package (v1.38.0) (10) and trimmed to the columns between *H. sapiens* residues 1176-1226. For each protein sequence, the number of non-gap characters was counted and plotted.

**ERC network analysis and visualization**

ERC network for TEFs was generated as described previously with limited modifications (11). Briefly, a Z-score cut-off of 3.5 was applied to identify factors that share a high ERC with each TEF (*PAF1* = 7, *CTR9* = 11, CDC73 = 7, *RTF1* = 26, *LEO1* = 7, *SPT4* = 12, *SPT5* = 17, *SPT6* = 33, *SPN1* = 3, *ELF1* = 4 genes passing threshold). Functions of genes were manually annotated using the Saccharomyces Genome Database (SGD) (12).

To verify that this TEF ERC network represents relevant connections and was not random, the global clustering coefficient of this network was compared to that of 10,000 randomly sampled networks. Each sampled network contained one random gene corresponding to each TEF to act as a query node. The top N genes with the highest ERC values with a query node were selected where N is the number of genes for the assigned TEF that passed the Z-score cutoff (e.g. the genes with the top 7 ERC values were collected for query nodes corresponding to *PAF1*). The resultant networks each had the same total number of nodes and the same number of edges originating from each query node as the TEF ERC network in **Fig. 5**.

**Software used for data visualization**

Sina plots in **Supplementary Fig. S4C** and **S11E** were made using Prism (v10.5.0). Other sina plots were generated using the geom_sina command in the ggforce package (v0.5.0) (13). UpSet plots were made in R using the ComplexUpSet package (v1.3.3) (14). Phylogenetic trees were visualized and annotated using iTOL (15). Networks in **Fig. 3C** and **Fig. 5** were generated in Cytoscape (v3.10.2) (16) and manually edited for visualization in Adobe Illustrator. Multiple sequence alignment snapshots were rendered using the AliView software (v1.28) (17).

**Data and code accessibility**

Code and data, including statistical test results, blacklisted proteins, RER and conservation score calculations, MSAs, ChimeraX sessions, AlphaFold predictions, final ortholog lists, summary of FoldSeek and PSI-BLAST search results, domain-specific HMMs, tree files, TEF ERC values are available within Zenodo repository **10.5281/zenodo.17856448**. See Zenodo README.md file in repository for a description of the repository layout.

**Supplementary Figures**

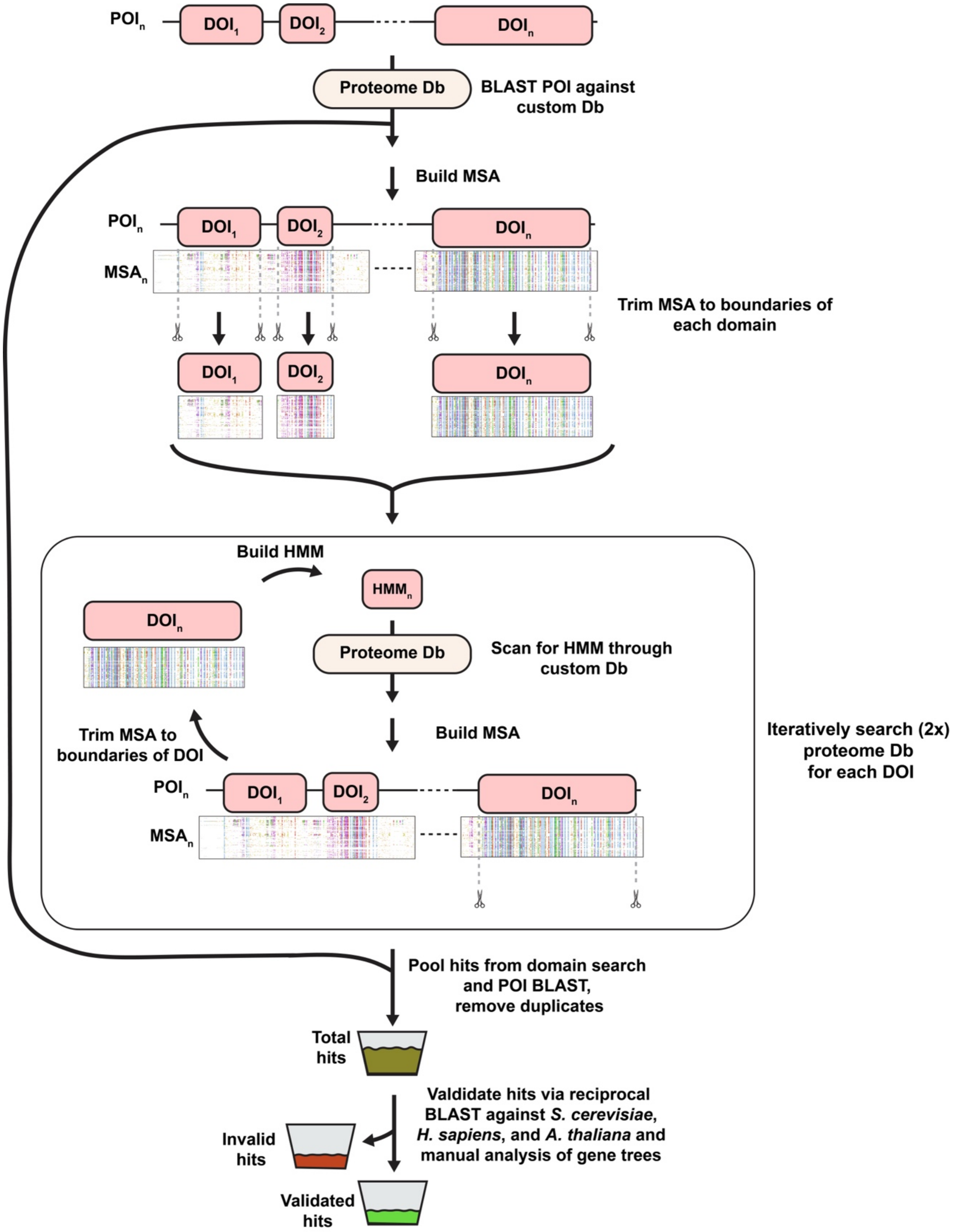


**Fig. S1. TEF homolog search strategy.** Diagram describing pipeline for TEF homolog searches combined from EukProt and GTDB databases. See Methods for more detail. POI- Protein of interest; DOI – Domain of interest; HMM – Hidden Markov Model; Db – Database; MSA – Multiple Sequence Alignment.

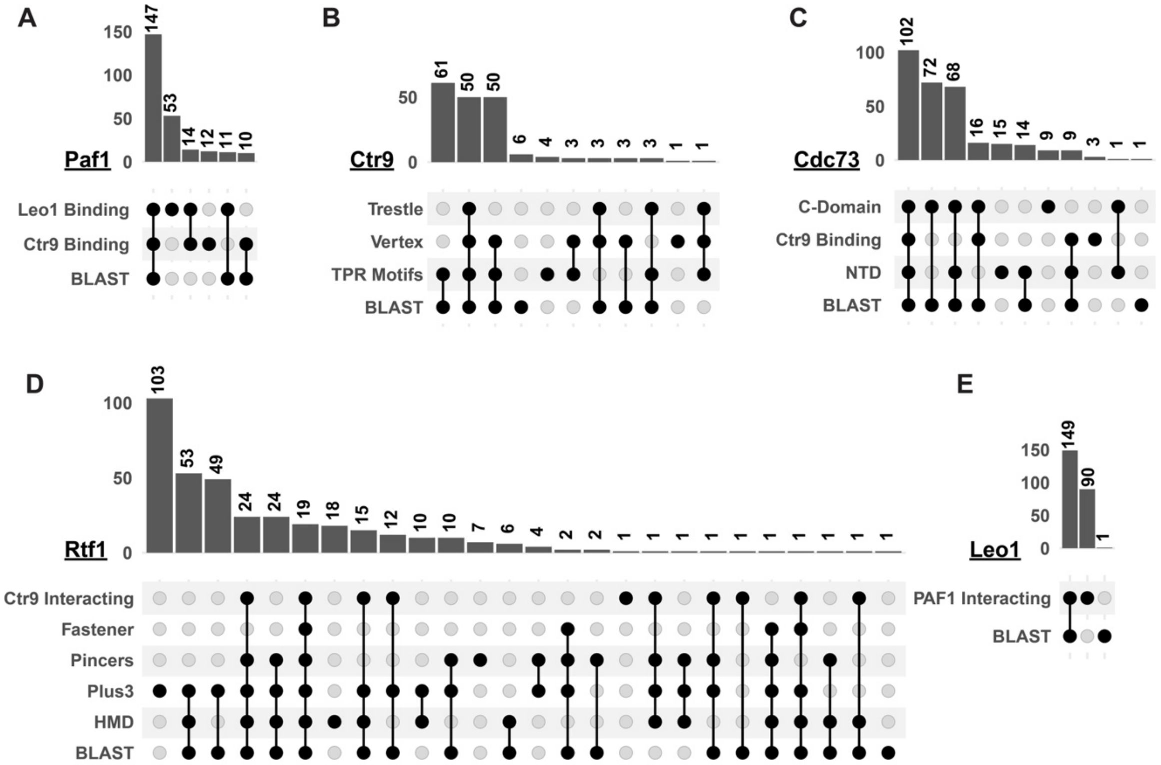


**Fig. S2. Comparison of HMM vs BLAST search efficacy for Paf1C subunits.** (A-E) UpSet plots depicting the number of homologs collected using BLAST and domain-specific HMM searches for (A) Paf1, (B) Ctr9, (C) Cdc73, (D) Rtf1, and (E) Leo1. Plots reflect the source of each protein hit, not the presence of each domain. For example, 53 Paf1 orthologs were identified by searching EukProt and GTDB proteomes using an HMM built from the Paf1 Leo1-binding domain but were not found when searching via the Paf1 Ctr9-binding domain or by BLAST with the human Paf1 sequence. This does not imply that in the 53 hits identified by searching for the Paf1 Leo1-binding domain, a Ctr9-binding domain was not detectable in the downstream analyses.

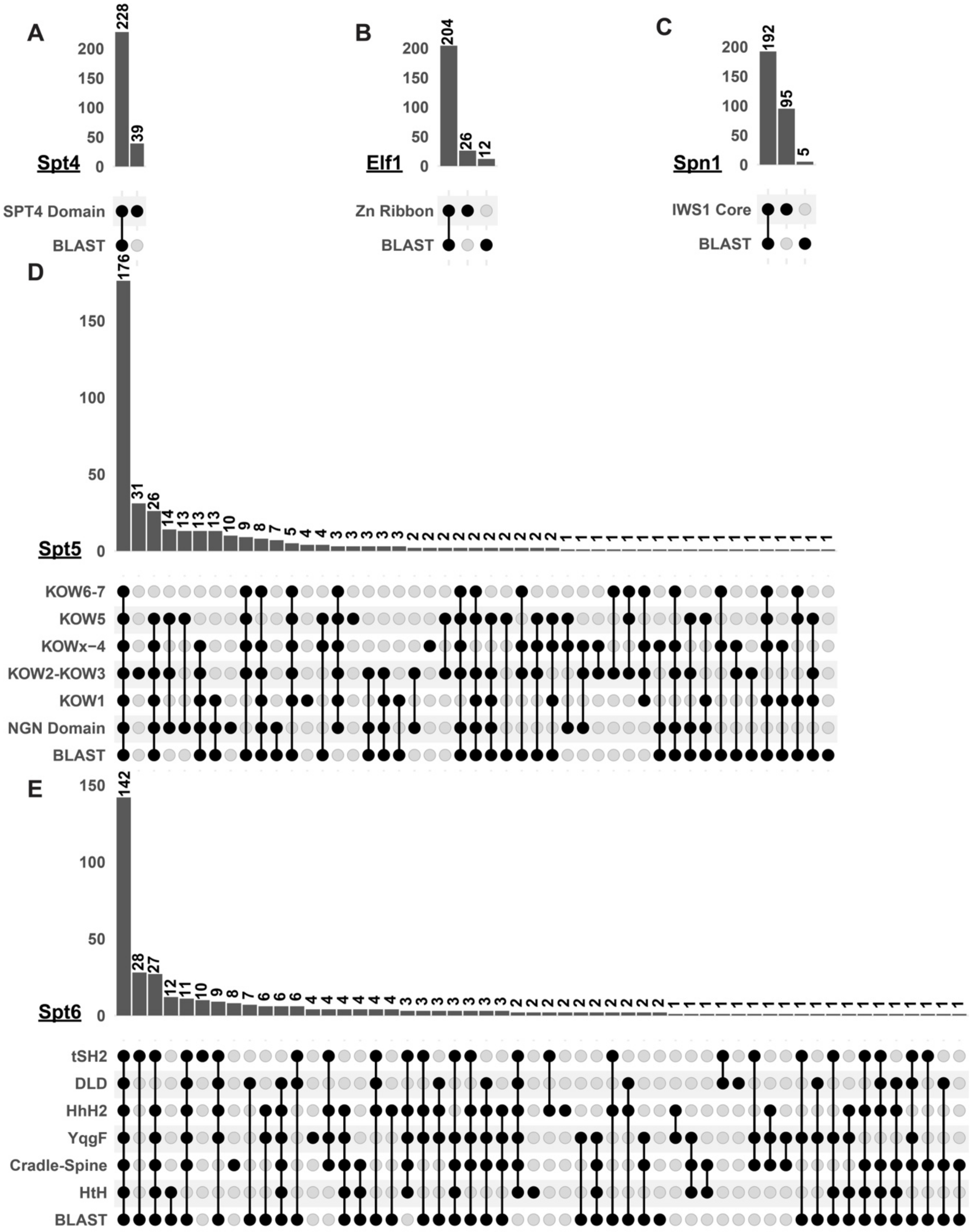


**Fig. S3. Comparison of HMM vs BLAST search efficacy for other TEFs.** (A-E) UpSet plots depicting the number of homologs collected using BLAST and domain-specific HMM searches for (A) Spt4, (B) Elf1, (C) Spn1, (D) Spt5, and (E) Spt6. Plots reflect the source of each protein hit, not the presence of each domain. See Fig. S2 for details.

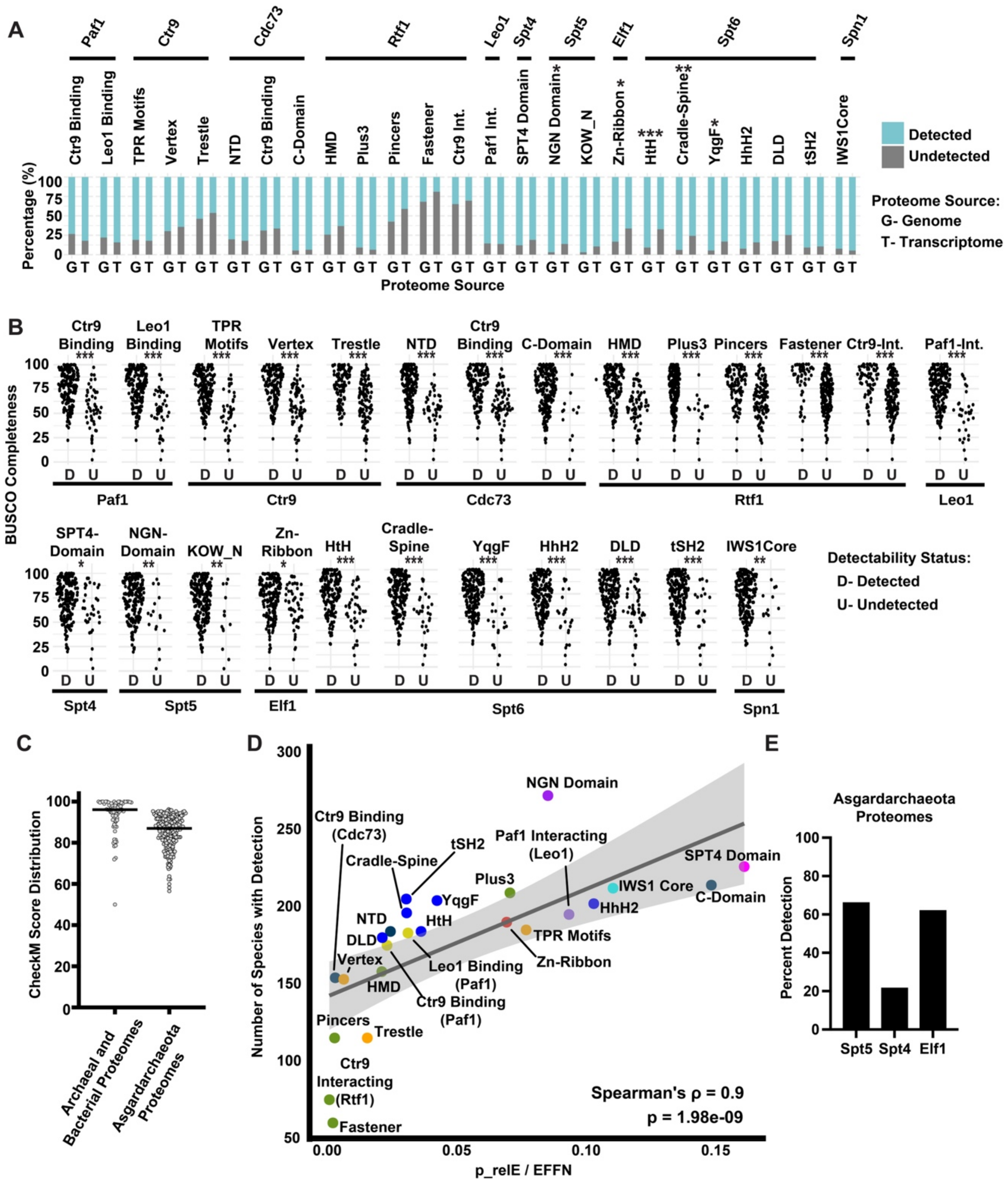


**Fig. S4. Detection of domains is dependent on the completeness of the source proteomes and the information content of the HMM.** (A) Stacked bar charts indicating the percentage of proteomes in which the domains were detected, grouped by the source of the predicted proteome. Fisher's Exact Test with multiple comparisons correction (Benjamini-Hochberg) was used to test if the probability of detecting a domain from genome-derived or transcriptome-derived proteomes is significantly different. (B) Sina plots comparing the distributions of BUSCO Completeness (18) of EukProt proteomes in which the indicated domain was detected versus not. Wilcoxon rank-sum test with multiple comparisons correction (Benjamini-Hochberg) was used to determine if the distribution of BUSCO proteome completeness scores is significantly correlated with domain detection across eukaryotic species. (C) Sina plot showing the distributions of CheckM completeness scores for GTDB proteomes used in this study. Left – Distribution of proteome scores used in the 304 species search. Right - Distribution of proteome scores used in the expanded Asgard archaea search. (D) Scatter plot depicting the relationship between the number of species in which a domain was detected and the information in the HMM. The ratio between p relE (mean positional relative entropy, in bits) and EFFN (effective sequence number) was used as a measure of HMM information content (19). Spearman's correlation was used to determine the relationship between the detection of the domain and information in the HMM. ρ-value indicates a positive correlation between the information in the HMM and detection of the domain. To further characterize the relationship between detectability of domains and the information in the HMMs, the data were fit to a linear regression model ($y = ß_0 + ß_1 \times \log(x)$). Band represents 95% confidence interval. (E) Bar plot indicating percentage of Asgardarchaeota species (n=218) in which Spt5, Spt4, and Elf1 orthologs were detected. See Methods for more details. *p < 0.05; **p < 0.01; ***p < 0.001.

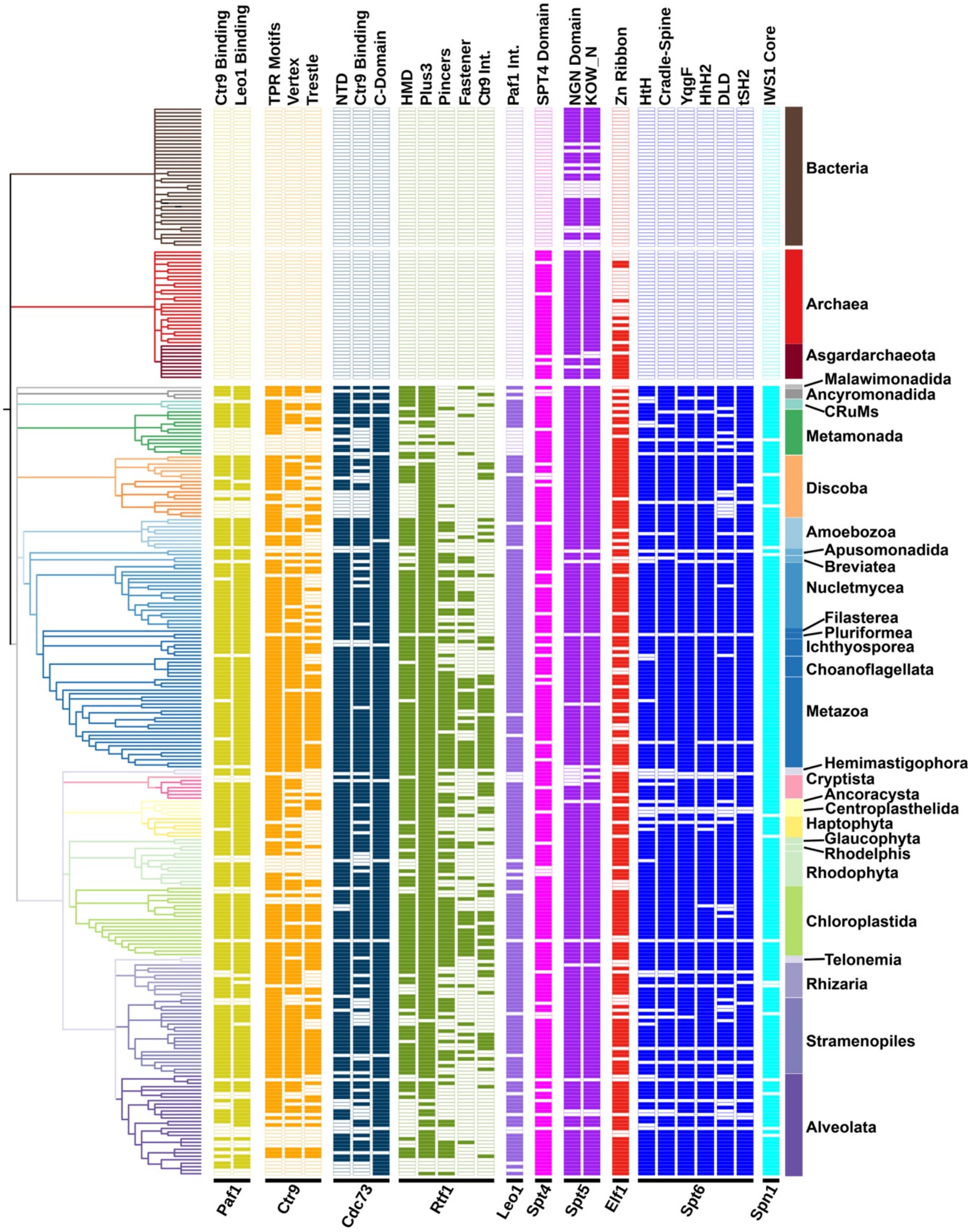


**Fig. S5. Species-level variation in detection of TEF domains.** Expanded tree from Fig. 1 depicting the detection status of TEF domains in each species in the combined EukProt/GTDB database. Each column corresponds to a domain in the indicated protein (listed at bottom), and each row corresponds to a proteome from an organism. A filled box indicates that the domain was detected in the proteome of the organism. Colored bars on the right represent clades.

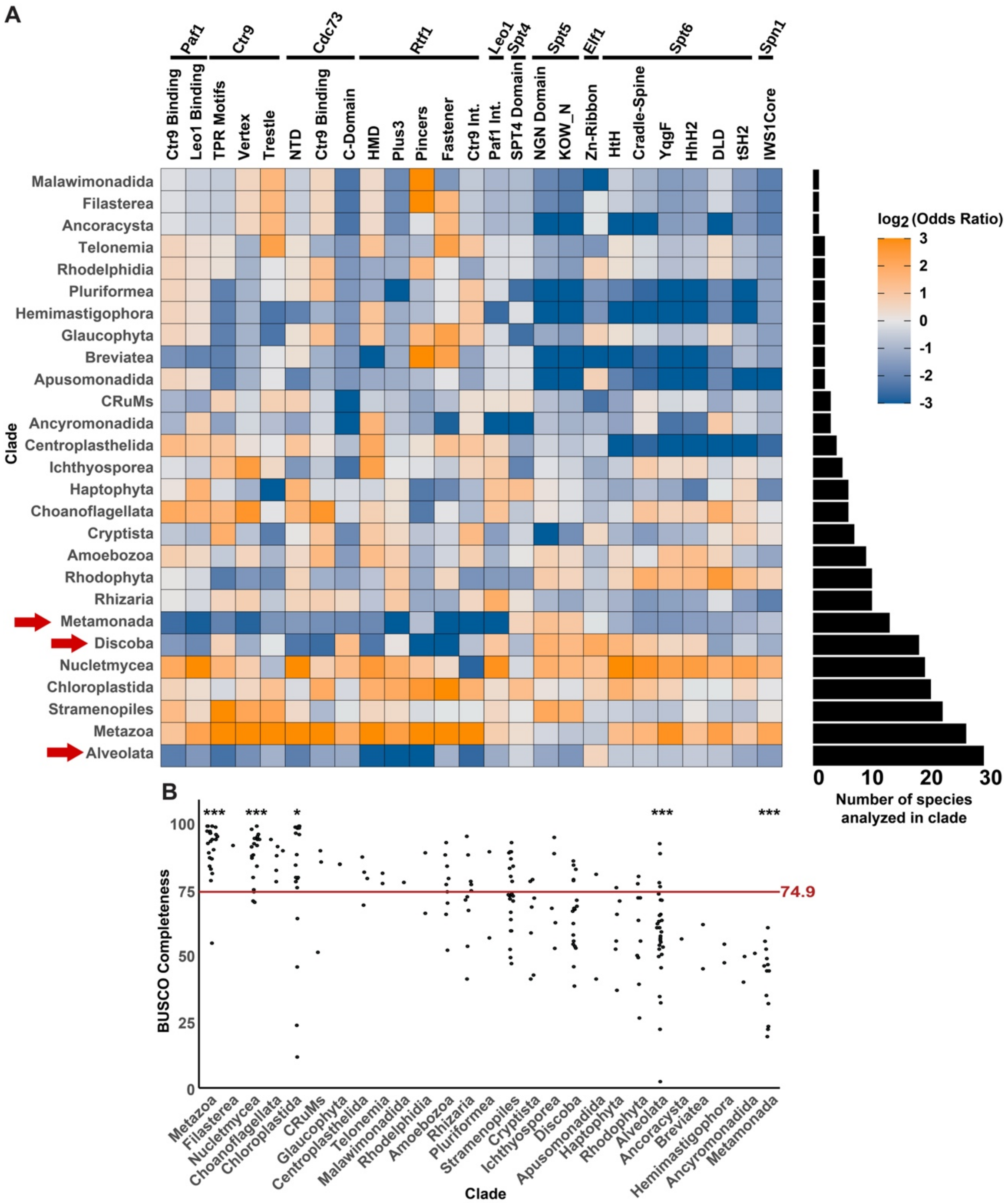


**Fig. S6. Odds of detecting Paf1C domains are lower in some clades.** (A) Heatmap showing $\log_2$(Odds Ratio) of domain detection in different clades. The Odds Ratio was calculated as follows: let C be a clade of interest and D be a domain of interest. Let x be the number of species in clade C in which domain D was detected. Let y be the number of species in which domain D was detected in clades other than clade C. Let $S_c$ be the total number of species in clade C. In total, our dataset contained proteomes from 227 eukaryotes. Therefore, for each domain-clade pair, an Odds Ratio was calculated as: $Odds\ Ratio = \frac{\frac{x}{S_C - x}}{\frac{y}{(227 - S_C) - y}}$. Bar plot on the right indicates number of species analyzed in each clade. Red arrows indicate clades in which the Odds Ratio for the detection of some Paf1C domains was less than 1. (B) Sina plots comparing the distributions of BUSCO completeness scores of EukProt proteomes from different clades. Red line and value indicate median BUSCO completeness score of all EukProt proteomes analyzed. Wilcoxon rank-sum test with multiple comparisons correction (Benjamini-Hochberg) was used to determine if the distribution of scores from proteomes in a clade significantly differ from the distribution of all proteomes in the database. $^{*}p < 0.05$; $^{**}p < 0.01$; $^{***}p < 0.001$.

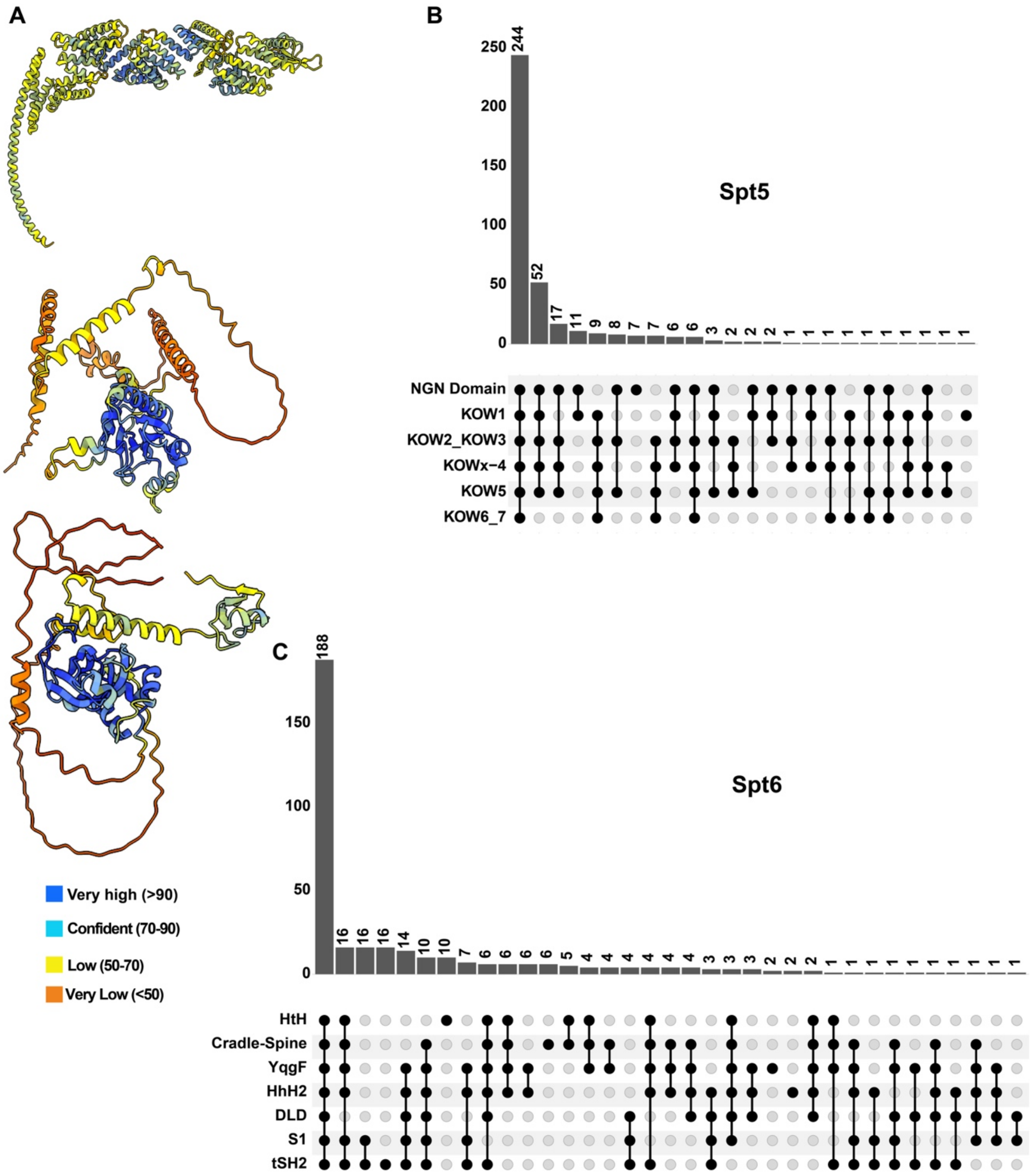


**Fig. S7. Multi-domain architectures of TEFs are broadly conserved.** (A) Models of AlphaFold3 predicted structures shown in Fig. 2, colored by the pLDDT score. Top – SsCtr9; middle - BsCdc73; bottom – NdRtf1. (B-C) UpSet plots depicting the coincidence of domain detection in homologs of indicated proteins as determined by HMMER hmmscan using custom HMMs. Scan domain eValue (--domE) threshold set to $10^{-3}$ (20).

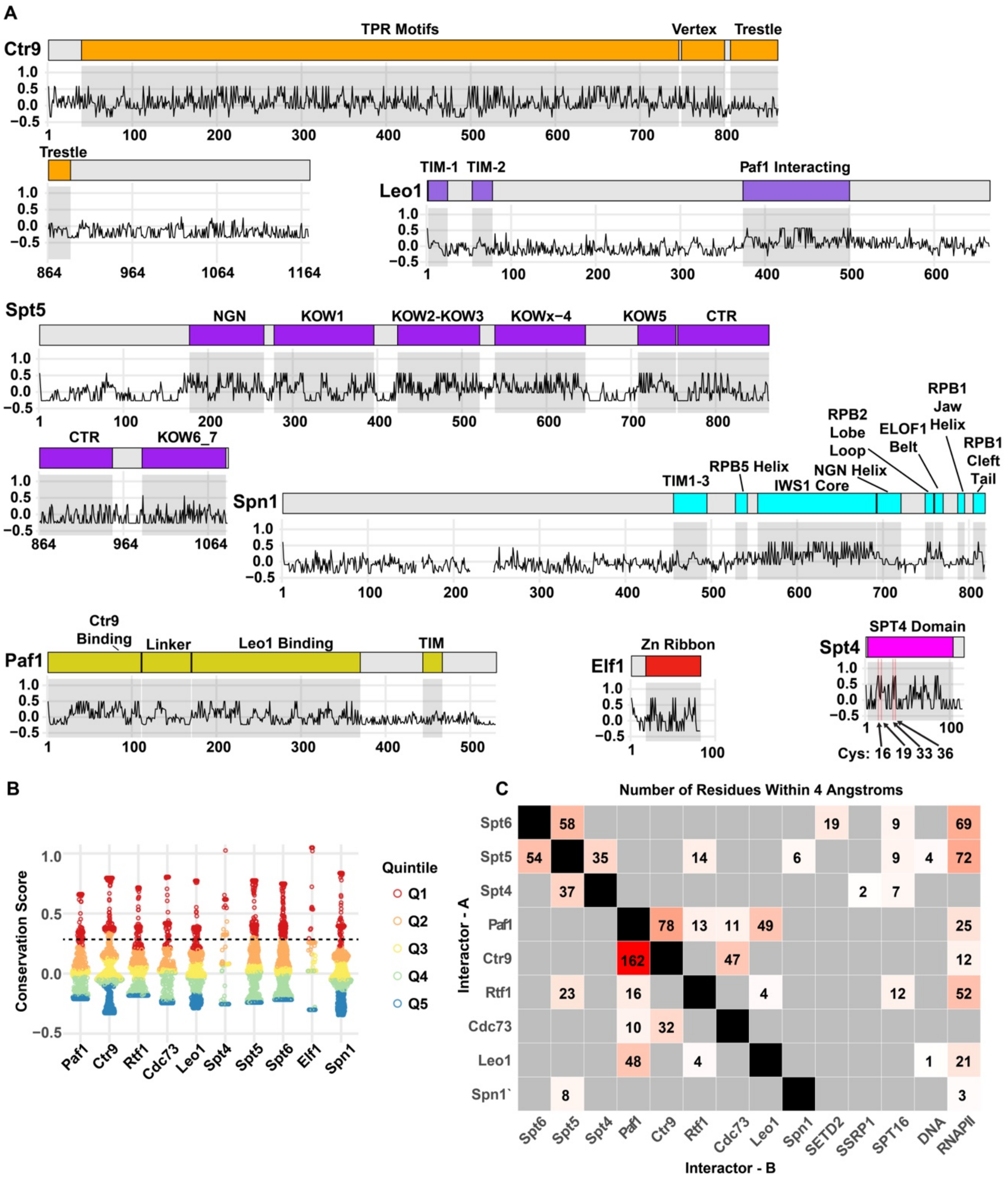


**Fig. S8. Conservation score analysis of TEF residues.** (A) Conservation scores of Ctr9, Leo1, Spt5, Spn1, Paf1, Elf1, and Spt4 across each residue along the *H. sapiens* homolog. See **Fig. 3** legend for a description of the plots. In the Spn1 line plot, gaps indicate residues that are unique to the human Spn1 homolog. (B) Sina plot showing the distribution of conservation scores of TEF residues. Data points have been colored by quintile. Dashed line indicates the cut-off used for residues considered as slowly evolving in **Fig. 3B**. (C) Diagram of pairwise interfaces in the transcription elongation complex (PDB: 9EH2). Number of residues of each TEF (y-axis/Interactor-A) within 4 angstroms of other components of the elongation complex (x-axis/Interactor-B). Color intensity scales with the number in each tile.

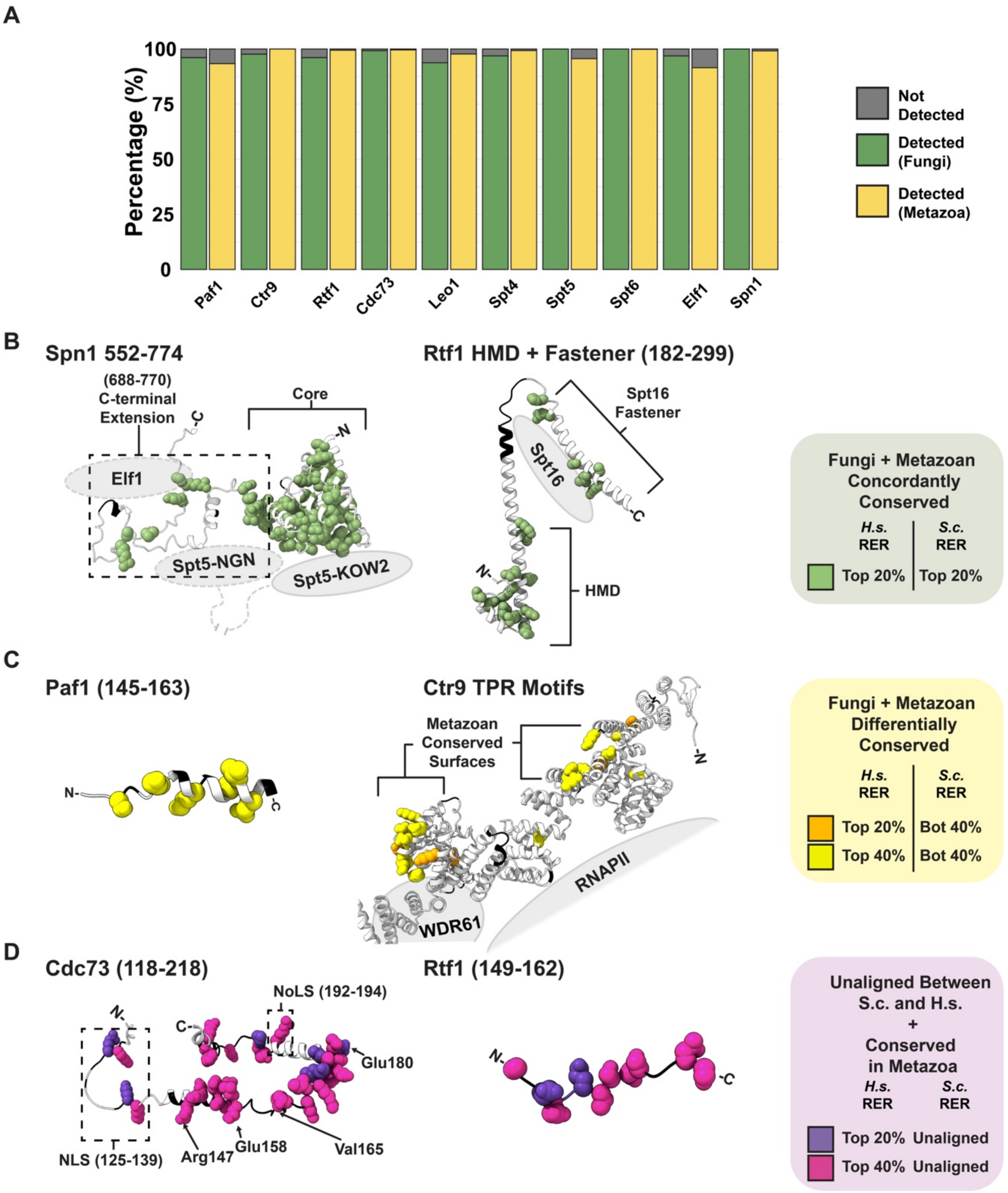


**Fig. S9. TEF detection and sequence conservation in metazoan and fungal proteome scans.** (A) Percentage of proteomes from RefSeq Fungi (n = 128) and RefSeq Metazoa (n = 706) databases for which a homolog was identified by BLAST search. (B-D) Different classes of conserved residues in TEFs, highlighted on the AlphaFold2 predicted structures of the human homologs. The relative positions of select additional factors on the transcription elongation complex are depicted as cartoon diagrams for clarity. Residues that are unaligned between the *S. cerevisiae* and *H. sapiens* orthologs and are not well conserved have been colored black. (B) Concordantly conserved residues (top 20% conserved residues in metazoan and fungal homologs) in the IWS1 domain of Spn1 (left) and the HMD of Rtf1 (right). Residues in Spn1 that are predicted to extend close to where Elf1 and Spt5 bind to elongating RNAPII are Arg751, Ala752, Val754, Tyr762, Arg765, and Pro766. Residues in Rtf1 that are conserved in both clades and interact with Spt16 are His263, Arg267, Ala281, Leu285, Ala287, and Arg289. (C) Differentially conserved residues (top 20-40% conserved residues in metazoan homologs and bottom 20% conserved residues in fungal homologs) in an uncharacterized region in Paf1 (left) and the TPR motifs of Ctr9 (right). (D) Top 20-40% conserved residues in metazoan homologs that are not mappable in the *S. cerevisiae* homolog, highlighted in understudied regions in Cdc73 (left) and Rtf1 (right).

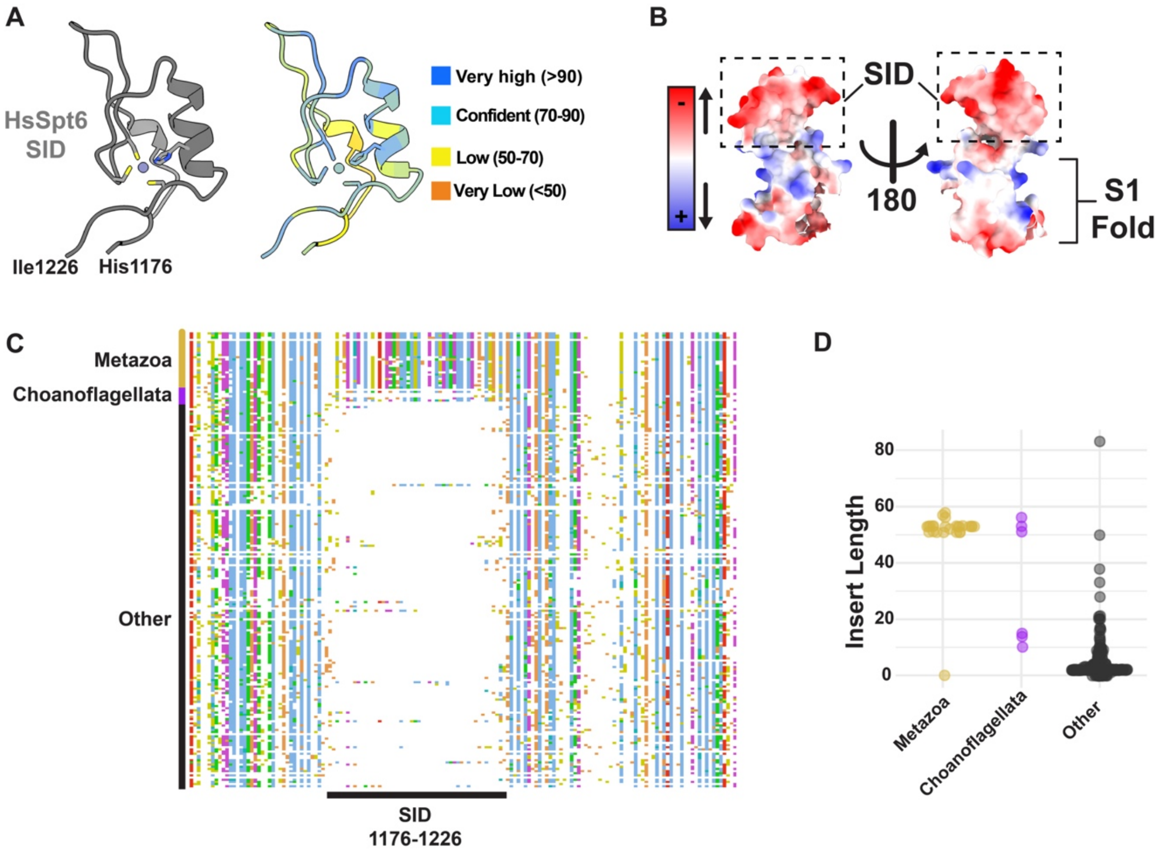


**Fig. S10. The S1 Insertion Domain is unique to metazoan and choanoflagellate Spt6 and is predicted to adopt a zinc-finger fold.** (A) Left - AlphaFold3 prediction of the human Spt6 S1 insertion domain (SID) with $Zn^{2+}$. Residues shown as sticks are predicted to coordinate a zinc ion. Right – AlphaFold3 prediction colored by pLDDT score. (B) Charge distribution of the human Spt6 S1 domain and SID surface residues displayed using the ChimeraX 'coulombic' command. (C) MSA snapshot of top-scoring Spt6 orthologs from eukaryotes, highlighting the insertion in Spt6 homologs in Metazoa and Choanoflagellata. MSA was trimmed using clipkit (--kpi-gappy) for clarity. (D) Sina plots showing the distribution of lengths of inserts in the S1 domain in Metazoa, Choanogflagellata, and other eukaryotic clades. MSA in panel C was examined to count the number of residues between positions aligning to the start and stop of the *H. sapiens* SID (residues 1176-1226). See Methods for more details.

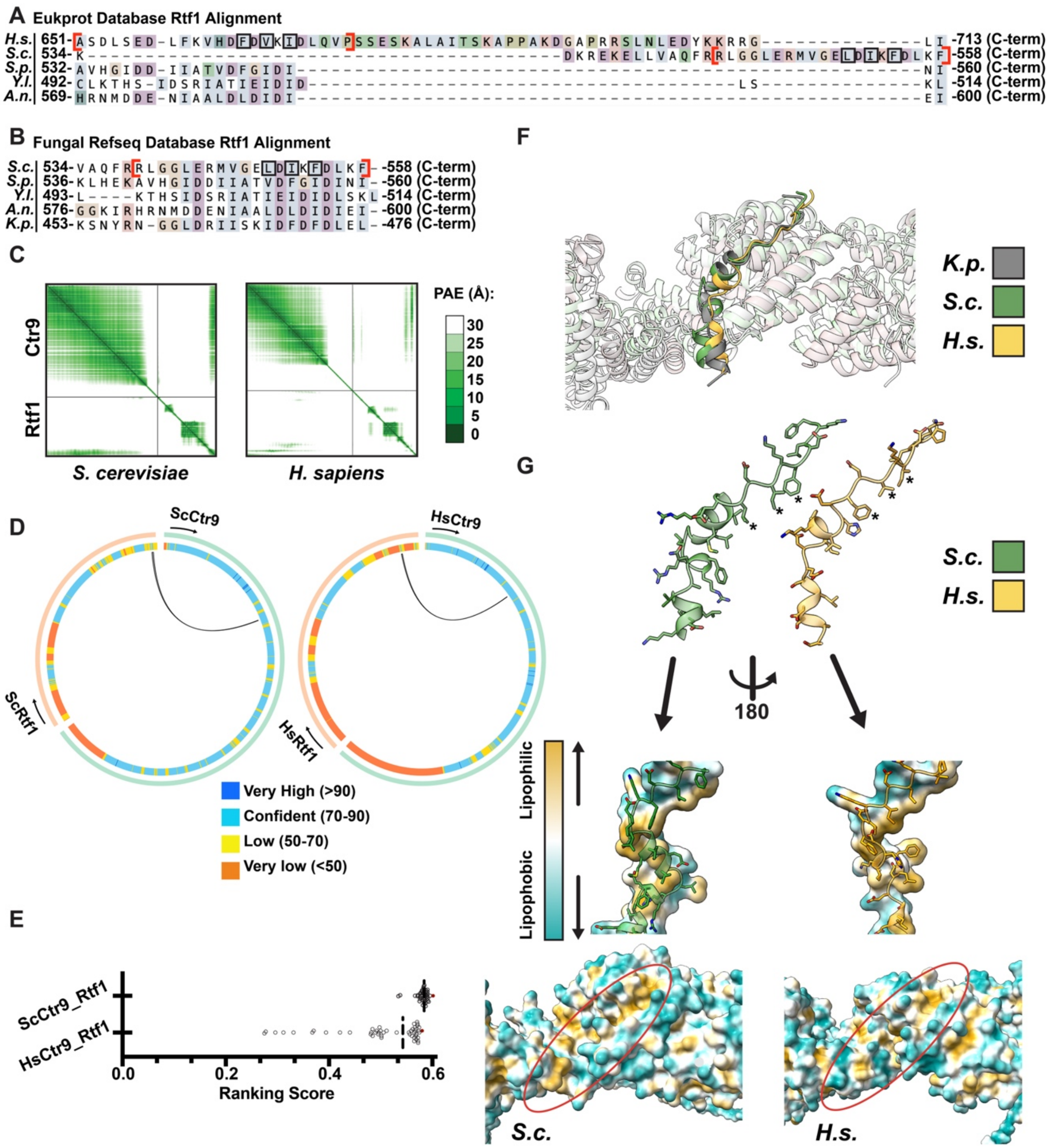


**Fig. S11. Human Rtf1 harbors a region homologous to fungal Rtf1 Hook.**
(A-B) MSA of Rtf1 homologs from EukProt/GTDB (A) and RefSeq fungal proteome (B) searches. Boundaries of the Rtf1 Hook are bracketed in red. Hydrophobic residues predicted to interact with a lipophilic surface in Ctr9 are labeled with black boxes. Regions corresponding to Rtf1 hook from the indicated species were collected, and all gap columns were removed. (A) Alignment of Rtf1 Hook region from *Homo sapiens*, *Saccharomyces cerevisiae*, *Schizosaccharomyces pombe*, *Yarrowia lipolytica*, and *Aspergillus nudilans*. (B) Alignment of Rtf1 Hook region from *Saccharomyces cerevisiae*, *Schizosaccharomyces pombe*, *Yarrowia lipolytica*, *Aspergillus nudilans*, and *Komagatella phaffi*. (C) Predicted aligned error (PAE) plots of top *S. cerevisiae* and *H. sapiens* Ctr9-Rtf1 co-fold predictions. PAE plot images were made in ChimeraX. (D) AlphaBridge plots for *S. cerevisiae* (top) and *H. sapiens* (bottom) Ctr9-Rtf1 co-folds, highlighting regions involved in interaction (black lines) between the two proteins. Inner rings represent the

pLDDT score of each residue. Predictions for *S. cerevisiae* and *H. sapiens* Ctr9-Rtf1 were generated using the AlphaFold3 web server. The output generated from the web server was downloaded as a .zip file and uploaded to the AlphaBridge web server (default parameters) to generate figures. (E) Distribution of AlphaFold3 Ranking Scores (n=50) for indicated protein pairs. Dotted line represents median of the distribution, and the brown point indicates the top-scoring model. The analysis in the following panels was done using this top-scoring model. (F) Rtf1 Hook in complex with Ctr9 as predicted by AlphaFold3 for *H. sapiens* (yellow) and *S. cerevisiae* (green) homologs, aligned to homologous cryo-EM structure of *Komagatella phaffi* proteins (PDB: 7XSX). (G) Lipophilicity maps of *H. sapiens* and *S. cerevisiae* Rtf1 Hook and Ctr9 grooves as calculated using the ChimeraX molecular lipophilicity potential (mlp) command. Black asterixis indicate hydrophobic residues highlighted in panels A and B.

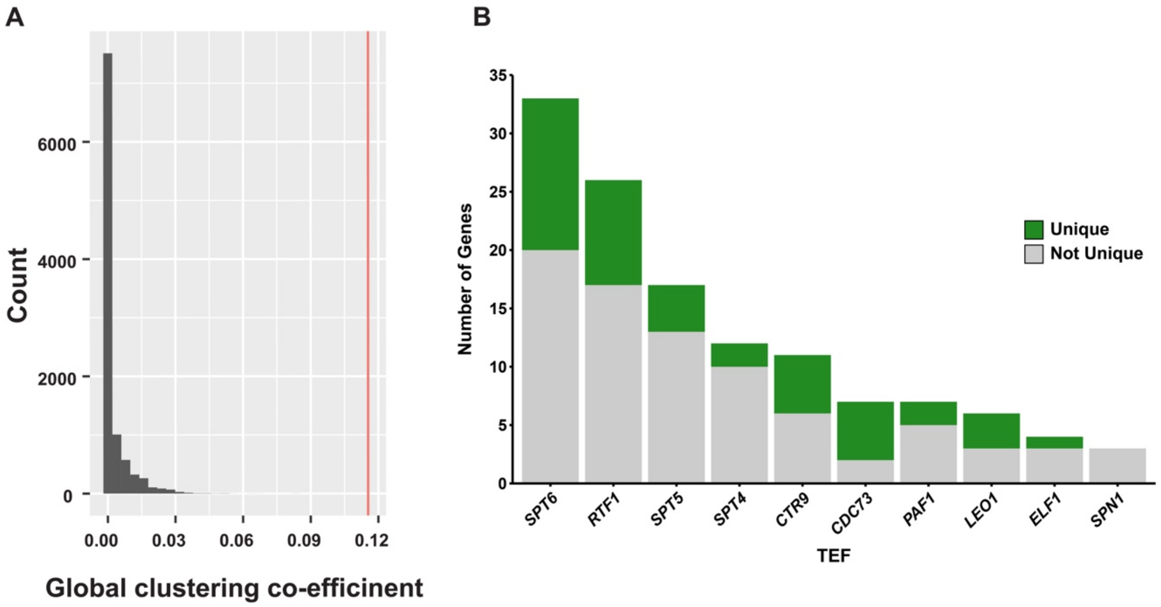


**Fig. S12. Global clustering coefficient of the TEF ERC network is greater than that of 10,000 randomly generated networks.** (A) Histogram showing the distribution of the global clustering coefficients of the 10,000 randomly sampled networks (see **Methods**). The red line represents the coefficient of the ERC network in **Fig. 5** (0.115). (B) Stacked bar plot showing the number of genes above Z-score ≥ 3.5 threshold connected to each TEF in the ERC network. Green bars represent the number of unique genes connected to the TEF in the network.

### Supplementary Tables

#### Table S1. Summary of alternative searches for domains in Discoba, Metamonada, and Alveolata.

| Protein | Domain | Clade | Sub-Clade_1 | Sub-Clade_2 | HMM Pipeline | FoldSeek | AlphaFold3 Prediction | PSI-BLAST |
|---|---|---|---|---|---|---|---|---|
| Paf1 | Leo1 Binding | Metamonada | Preaxostyla | - | Yes | - | | |
| | | | Parabasalia | - | No | Inconclusive[a] | No Hit | No |
| | | | Anaeramoebidae | - | Yes | - | | |
| | | | Fornicata | - | No | Inconclusive[a] | No Hit | No |
| Ctr9 | TRP Motifs | | Preaxostyla | - | Yes | - | | |
| | | | Parabasalia | - | Yes | | | |
| | | | Anaeramoebidae | - | Yes | | | |
| | | | Fornicata | - | No | Inconclusive[a] | No Hit | No |
| | Vertex | | Preaxostyla | - | Yes | - | | |
| | | | Parabasalia | - | No | No | Yes | - |
| | | | Anaeramoebidae | - | Yes | - | | |
| | | | Fornicata | - | No | No | No Hit | No |
| | Trestle | | Preaxostyla | - | Yes | - | | |
| | | | Parabasalia | - | Yes | | | |
| | | | Anaeramoebidae | - | Yes | | | |
| | | | Fornicata | - | No | No | No Hit | No |
| Rtf1 | HMD | | Preaxostyla | - | Yes | - | | |
| | | | Parabasalia | - | No | No | No Hit | No |
| | | | Anaeramoebidae | - | Yes | - | | |
| | | | Fornicata | - | Yes | | | |
| | Plus3 | | Preaxostyla | - | Yes | - | | |
| | | | Parabasalia | - | No | No | No Hit | No |
| | | | Anaeramoebidae | - | Yes | - | | |
| | | | Fornicata | - | Yes | | | |
| Leo1 | Paf1 Binding | | Preaxostyla | - | Yes | - | | |
| | | | Parabasalia | - | No | Inconclusive[a] | No Hit | No |
| | | | Anaeramoebidae | - | Yes | - | | |
| | | | Fornicata | - | No | Inconclusive[a] | No Hit | No |
| Paf1 | Leo1 Binding | Discoba | Jakobida | - | Yes | - | | |
| | | | Heterolobosea | - | Yes | | | |
| | | | Euglenozoa | Kinetoplastea | No | Inconclusive[a] | No Hit | No |
| | | | | Diplonemea | Yes | - | | |
| | | | | Euglenida | Yes | | | |
| Ctr9 | Vertex | | Jakobida | - | Yes | - | | |
| | | | Heterolobosea | - | Yes | | | |
| | | | Euglenozoa | Kinetoplastea | Yes | | | |
| | | | | Diplonemea | No | No | Yes | - |
| | | | | Euglenida | Yes | - | | |
| Cdc73 | NTD | | Jakobida | - | Yes | - | | |
| | | | Heterolobosea | - | Yes | | | |
| | | | Euglenozoa | Kinetoplastea | No | No | No | No |
| | | | | Diplonemea | No | No | No | No |
| | | | | Euglenida | Yes | - | | |
| Rtf1 | HMD | | Jakobida | - | Yes | - | | |
| | | | Heterolobosea | - | Yes | | | |
| | | | Euglenozoa | Kinetoplastea | No | No | No | No |
| | | | | Diplonemea | No | No | No | No |
| | | | | Euglenida | Yes | - | | |
| Cdc73 | NTD | Alveolata | Colponemidae | - | Yes | - | | |
| | | | Ciliophora | - | Yes | | | |
| | | | Myzozoa | Dinoflagellates | Yes | | | |
| | | | | Colpodellida | No | Yes | - | |
| | | | | Apicomplexa | Yes | - | | |
| Rtf1 | HMD | | Colponemidae | - | Yes | - | | |
| | | | Ciliophora | - | Yes | | | |
| | | | Myzozoa | Dinoflagellates | No | No | No | No |
| | | | | Colpodellida | No | Yes | - | |
| | | | | Apicomplexa | Yes | - | | |

[a]Since Paf1 and Leo1 domains form beta-barrel structures, their FoldSeek searches were confounded by other proteins that adopt a similar fold. As such, FoldSeek searches for these domains remained inconclusive.

### References


1. M. van Kempen *et al.*, Fast and accurate protein structure search with Foldseek. *Nat Biotechnol* **42**, 243-246 (2024).
2. S. B. Van Oss *et al.*, The Histone Modification Domain of Paf1 Complex Subunit Rtf1 Directly Stimulates H2B Ubiquitylation through an Interaction with Rad6. *Mol Cell* **64**, 815-825 (2016).
3. W. Sun *et al.*, Crystal structure of the N-terminal domain of human CDC73 and its implications for the hyperparathyroidism-jaw tumor (HPT-JT) syndrome. *Sci Rep* **7**, 15638 (2017).
4. P. Deng *et al.*, Transcriptional elongation factor Paf1 core complex adopts a spirally wrapped solenoidal topology. *Proc Natl Acad Sci U S A* **115**, 9998-10003 (2018).
5. A. D. Wier, M. K. Mayekar, A. Heroux, K. M. Arndt, A. P. VanDemark, Structural basis for Spt5-mediated recruitment of the Paf1 complex to chromatin. *Proc Natl Acad Sci U S A* **110**, 17290-17295 (2013).
6. J. Abramson *et al.*, Accurate structure prediction of biomolecular interactions with AlphaFold 3. *Nature* **630**, 493-500 (2024).
7. S. F. Altschul *et al.*, Gapped BLAST and PSI-BLAST: a new generation of protein database search programs. *Nucleic Acids Res* **25**, 3389-3402 (1997).
8. K. Tamura, G. Stecher, S. Kumar, MEGA11: Molecular Evolutionary Genetics Analysis Version 11. *Mol Biol Evol* **38**, 3022-3027 (2021).
9. E. C. Meng *et al.*, UCSF ChimeraX: Tools for structure building and analysis. *Protein Sci* **32**, e4792 (2023).
10. J. D. Thompson, D. G. Higgins, T. J. Gibson, CLUSTAL W: improving the sensitivity of progressive multiple sequence alignment through sequence weighting, position-specific gap penalties and weight matrix choice. *Nucleic Acids Res* **22**, 4673-4680 (1994).
11. J. H. Little *et al.*, ERC2.0 evolutionary rate covariation update improves inference of functional interactions across large phylogenies. *Genome Res* 10.1101/gr.280586.125 (2025).
12. E. D. Wong *et al.*, *Saccharomyces* genome database update: server architecture, pan-genome nomenclature, and external resources. *Genetics* **224** (2023).
13. T. Pedersen L. (2025) ggforce: Accelerating 'ggplot2'.
14. M. Krassowski (2021) ComplexUpset: Create Complex UpSet Plots Using 'ggplot2' Components.
15. I. Letunic, P. Bork, Interactive Tree of Life (iTOL) v6: recent updates to the phylogenetic tree display and annotation tool. *Nucleic Acids Res* **52**, W78-W82 (2024).
16. P. Shannon *et al.*, Cytoscape: a software environment for integrated models of biomolecular interaction networks. *Genome Res* **13**, 2498-2504 (2003).
17. A. Larsson, AliView: a fast and lightweight alignment viewer and editor for large datasets. *Bioinformatics* **30**, 3276-3278 (2014).
18. F. A. Simao, R. M. Waterhouse, P. Ioannidis, E. V. Kriventseva, E. M. Zdobnov, BUSCO: assessing genome assembly and annotation completeness with single-copy orthologs. *Bioinformatics* **31**, 3210-3212 (2015).
19. S. R. Eddy, Accelerated Profile HMM Searches. *PLoS Comput Biol* **7**, e1002195 (2011).
20. S. C. Potter *et al.*, HMMER web server: 2018 update. *Nucleic Acids Res* **46**, W200-W204 (2018).
